# Deleterious impacts of inactivity and beneficial impacts of neuromuscular electrical stimulation on muscle structure and function in the zebrafish model of Duchenne Muscular Dystrophy

**DOI:** 10.1101/2020.09.02.279513

**Authors:** Elisabeth A. Kilroy, Kaylee L. Brann, Claire E. Schaffer, Devon Varney, Kodey J. Silknitter, Jordan N. Miner, Ahmed Almaghasilah, Tashawna L. Spellen, Alexandra D. Lewis, Karissa Tilbury, Benjamin L. King, Joshua B. Kelley, Clarissa A. Henry

## Abstract

Although it is known that inactivity is deleterious for healthy individuals, less is known about the consequences of inactivity on muscle disease. Reduced activity is frequently encouraged for individuals with congenital muscular dystrophies such as Duchenne Muscular Dystrophy (DMD). We used the zebrafish *dmd* mutant and a longitudinal design to elucidate the consequences of inactivity versus activity on muscle health. Inactivity worsened muscle structure and survival. We designed four neuromuscular stimulation paradigms loosely based on weight lifting regimens. Each paradigm differentially affected muscle structure, function, and survival. Only endurance neuromuscular stimulation (eNMES) improved all outcome measures. We found that eNMES (1) returns gene expression to wild-type levels, (2) increases muscle adhesion to the extracellular matrix (ECM), and (3) remodels the ECM and supports regeneration. Our data indicate that inactivity is deleterious but neuromuscular stimulation can be beneficial, suggesting that the right type of activity may benefit patients with muscle disease.

## INTRODUCTION

Skeletal muscle is a dynamic tissue whose structural and molecular networks change in response to demand. Skeletal muscle’s ability to adapt is critical to maintaining not only muscle health, but also the overall health of an individual. Skeletal muscle is one of the primary predictors of longevity and recovery from illness and injury, demonstrating that robust skeletal muscle mass is essential for whole-body homeostasis (Margolis and Rivas, 2015). Whereas a great deal is known about the structural and functional plasticity of healthy skeletal muscle, far less is understood about plasticity and adaptation in diseased muscle. Congenital muscular dystrophies are debilitating progressive diseases without cures. It seems important to learn how inactivity and activity impact muscle wasting in congenital muscular dystrophies. Here, we describe how inactivity and neuromuscular electrical stimulation (NMES) impact the zebrafish model of Duchenne Muscular Dystrophy (DMD).

Individuals with DMD harbor mutations in the gene encoding the protein dystrophin (Hoffman et al., 1987). Dystrophin provides a link between the actin cytoskeleton and the extracellular matrix (ECM) and serves as a scaffold for the assembly of the dystrophin-glycoprotein complex (DGC) within the sarcolemma (muscle plasma membrane) (Bonilla et al., 1988; Ervasti and Campbell, 1991). The stability and integrity of the ECM and DGC are critical to the viability of muscle fibers during contraction. The ECM modulates mechanical homeostasis and cell-matrix interactions (Grzelkowska-Kowalczyk, 2016; Humphrey et al., 2014). ECM-mediated distribution and transmission of force across muscle fibers is mostly achieved through the ECM-cytoskeleton linkage via the DGC (Humphrey et al., 2014; Ramaswamy et al., 2011). The DGC maintains the structural integrity of the sarcolemma and serves as a scaffold for various signaling and channel proteins as well as an anchoring point for signaling molecules near their sites of action (Constantin, 2014). The progressive muscle wasting and weakness in DMD is thought to result at least in part because the lack of dystrophin and the disruption of the DGC mechanically weakens the sarcolemma. Stress placed on a mechanically weakened sarcolemma causes microlesions to develop along the sarcolemma, increasing calcium entry, which results in muscle protein degradation and muscle fiber necrosis (Alderton and Steinhardt, 2000; Gailly, 2002; Gillis, 1996; Ruegg et al., 2002). Given the role that the DGC plays in muscle during force generation and the concern about potentially increased muscle damage, there has been interest in studying the effects of activity on the progression of muscle disease. However, studies tend to be small with heterogeneous populations and a clear answer has not yet been established (Alemdaroğlu et al., 2015; Bushby et al., 2010; Gianola et al., 2013; Hyzewicz et al., 2015; Jansen et al., 2013; Markert et al., 2012). It is well-established that individuals with DMD exhibit lower activity levels compared to their healthy counterparts beginning at a very young age (Jeannet et al., 2011; McDonald, 2002), yet the consequences of these reduced activity levels on disease progression are not fully understood. Reduced activity is associated with worsening muscle quality and mass in healthy individuals. This raises the question of how inactivity could affect DMD, especially because lower-limb strength and gait velocity are the most important predictors of functional ambulation and independence in individuals with DMD (McDonald et al., 1995).

Skeletal muscle fibers are influenced by the activity pattern imposed upon them, whether via the innervating neuron or electrical stimulation (Pette and Vrbová, 1985). Early researchers, including Guillaume Benjamin Amand Duchenne, the French neurologist who described DMD in 1861, proposed that super-imposing electrical stimulation on dystrophin-deficient muscles could serve as a potential therapy (Barnard et al., 1986; Duchenne, GB, 1870; Reichmann et al., 1981). NMES delivers a series of waveforms of electrical current that is characterized by its frequency, amplitude, and pulse width (or pulse duration) (Sheffler and Chae, 2007). These three parameters dictate the strength of the muscle contraction and the amount of force that is generated. The main advantage of NMES is its ability to activate fast- and slow-twitch muscle fibers resulting in hypertrophy without high-effort voluntary force generation (Gondin et al., 2011). It is possible that NMES could impact muscle deterioration in congenital muscular dystrophies. However, only a few studies have examined the impact of NMES on muscle strength and function in humans (Scott et al., 1990, 1986; Zupan, 1992; Zupan et al., 1993) and the *mdx* mouse model of DMD (Dangain and Vrbova, 1989; Luthert et al., 1980; Vrbová and Ward, 1981). The only clear conclusion from these studies is that NMES does not appear to be detrimental.

Zebrafish are an attractive model to elucidate the consequences of inactivity versus NMES on diseased muscle. Many molecular, ultrastructural and histological features are shared between zebrafish and human muscle, including components of the DGC, the excitation-contraction coupling machinery, and the contractile apparatus (Dou et al., 2008; Dowling et al., 2009; Guyon et al., 2003; Parsons et al., 2002). Further, zebrafish exhibit reproducible, quantitative motor behaviors beginning at 1 dpf (Saint-Amant and Drapeau, 1998), providing simple and non-invasive measures of muscle function. Dystrophin-deficient zebrafish, known as sapje^ta222a/ta222a^, and referred here as *dmd* mutants, are the smallest vertebrate model of DMD. These *dmd* mutants exhibit severe structural and functional deficits by 4 dpf, and die prematurely between their second and third weeks (Bassett, 2003; Berger et al., 2010).

The purpose of this study was to use a longitudinal study design to evaluate the impacts of inactivity and NMES on disease progression in *dmd* mutant zebrafish. This longitudinal design showed for the first time that there is consistent phenotypic variation in *dmd* zebrafish that impacts the progression of muscle degeneration. We found that inactivity lowered the threshold for contraction-induced injury and reduced lifespan. Interestingly, different NMES programs that varied in pulse frequency and voltage had differing effects on muscle structure, neuromuscular junction structure, motility, and lifespan. We did identify one program that improved all of the above. This program also increased resilience of muscle fibers to high-force contraction, increased sarcomere length, and improved nuclear morphology. Deep sequencing indicated that wild-type (WT) and *dmd* muscle responded differently to NMES, with the main effect of NMES being to shift gene expression closer to the WT profile. Taken together, our results show that NMES can have a dramatic positive impact on *dmd* muscle and establish the zebrafish as an excellent model for longitudinal studies that are critical for elucidating basic mechanisms of skeletal muscle plasticity.

## METHODS

### Zebrafish Husbandry and Transgenic Lines

Zebrafish embryos were retrieved from natural spawns of adult zebrafish maintained on a 14-h light/10-h dark cycle. We used sapje^ta222a^ zebrafish (Bassett, 2003). For live imaging studies, we used transgenic sapje^ta222a^ 3MuscleGlow zebrafish expressing mylpfa:lyn-cyan, smych1:GFP, and myog:H2B:RFP (gift from Drs. Sharon Amacher and Jared Talbot) (Hromowyk et al., 2020). Embryos were grown in embryo rearing media (ERM) with methylene blue at 28.5 degrees Celsius. Embryos were manually dechorionated at 1 dpf. Zebrafish were fed once daily beginning at 5 dpf (Larval AP100 Dry Larval Diet (<50 microns), Zeigler, Pennsylvania, USA). For survival studies, zebrafish were housed in 20 mm petri dishes with 10 mL of system water per dish beginning at 8 dpf. Survival checks were performed in the morning and at night. All protocols conform to the University of Maine Institutional Animal Care and Use Committee’s Guidelines.

### Experimental Overview

Experiments were conducted identically so that variables such as treatment duration, disease stage at time of treatment, and disease stage at time of evaluation did not change. Zebrafish were followed individually throughout each experiment so that disease progression could be monitored throughout time in longitudinal studies. Experiments begin at disease onset. Disease onset for our sapje line is at 2 dpf. For the live imaging studies, disease onset in the transgenic sapje line is at 3 dpf even though the alleles harboring the mutation are identical. However, this transgenic line was imported from The Ohio State University, while our line has been maintained solely at University of Maine. Experiments were carried out exactly the same for our sapje line and the transgenic line. At disease onset, zebrafish were identified via birefringence as a *dmd* mutant or healthy wild-type (WT) sibling. Healthy WT siblings had myotomes with organized, parallel muscle fibers that appear bright white while *dmd* mutants had myotomes with disorganized and detached muscle fibers that appear gray to black (Bassett, 2003). Larvae were housed in 12-well plates (one fish per well) with 3 mL of ERM per well. Zebrafish were then randomly assigned to control, inactivity, or NMES cohorts for the next three days (the treatment period). At the end of this treatment period, zebrafish were allowed to recover for an additional four days (recovery period). During the treatment and recovery periods, disease progression was monitored by daily birefringence and swim activity analyses, with a special emphasis on what is occurring at 5 and 8 dpf as these mark the beginning and end of the recovery period.

### Inactivity paradigm

For intermittent inactivity, zebrafish were placed in a low dose of tricaine (MS-222, Sigma-Aldrich, Missouri, USA; 306 μM in 1X ERM) overnight for 12 hours each day for three days beginning at disease onset (2, 3, and 4 dpf). The total time that zebrafish were inactive was 36 hours (Figure 2A1). For extended inactivity, zebrafish were placed in the same dose of tricaine for 72 hours beginning at disease onset (2, 3, and 4 dpf), and were removed from tricaine at the start of 5 dpf (Figure 2A2). Birefringence images were taken at disease onset, immediately prior to removal from tricaine on 5 dpf, and three days following removal from tricaine (8 dpf). DanioVision was used to evaluate swim function in response to intermittent or extended inactivity, and was performed at 8 dpf. Following DanioVision, zebrafish were either fixed for further analyses of muscle health via immunostaining or followed daily for survival.

### NMES paradigm

The first NMES session began at disease onset. The treatment period included 1 session of NMES at 2, 3 and 4 dpf for a total of 3 sessions. Following the completion of the third NMES session, zebrafish entered the recovery period from 5 to 8 dpf.

Zebrafish were subjected to NMES in groups of four using our 3D printed ‘gym’ (Figure 4B). The rectangular gym is divided into 6 rectangular wells that measure 4.7625 mm (length), 1.5875 mm (width), and 1.5875 mm (depth). Two tunnels run parallel to the smaller sides of the rectangular wells and the positive and negative electrodes slide through these tunnels such that they are exposed only in the wells. This allows the delivery of electrical pulses to each zebrafish simultaneously. Prior to the NMES session, zebrafish were transferred to tricaine solution (612 μM in 1X ERM) for 4 minutes. At the end of the 4 minutes, zebrafish were placed into a well with its head facing the positive electrode and its tail facing the negative electrode. The positive and negative electrodes were attached to a Grass SD9 Stimulator, which was used to generate the electrical pulses. The frequency, delay, and voltage were adjusted for the different paradigms (Figure 4C and D). Each NMES session lasted 1 minute. Following each NMES session, zebrafish were removed from the gym and placed back into their respective wells.

### Birefringence Analysis

Birefringence was used to quantitatively assess the daily progression of dystrophy (Berger et al., 2012). Zebrafish were placed in tricaine (612 μM) immediately prior to imaging and then transferred to a 35-mm glass bottom dish. Birefringence images were taken on a Leica MZ10 F Stereomicroscope with a Zeiss AxioCam MRm or Leica DMC5400 camera attached. An analyzer in a rotatable mount (Leica) was attached to the objective and the glass-bottom petri dish was placed on the polarized glass stage. Images were taken at the same time every day within an experiment. Imaging parameters were consistent for all zebrafish and across all days. Mean gray values were calculated using FIJI software as described previously (Berger et al., 2012). Briefly, the body of the zebrafish was outlined from the 6th to the 25th myotome using the “Polygon selections” tool and then the mean gray value was measured. Three separate outlines were drawn to obtain three separate measures, and the average was used for calculations. All images were blinded prior to measurements using a Perl script. Mean gray values are presented as a percentage of the average mean gray value of healthy WT siblings in the control group (equation in Figure 1A).

**Figure 1:**
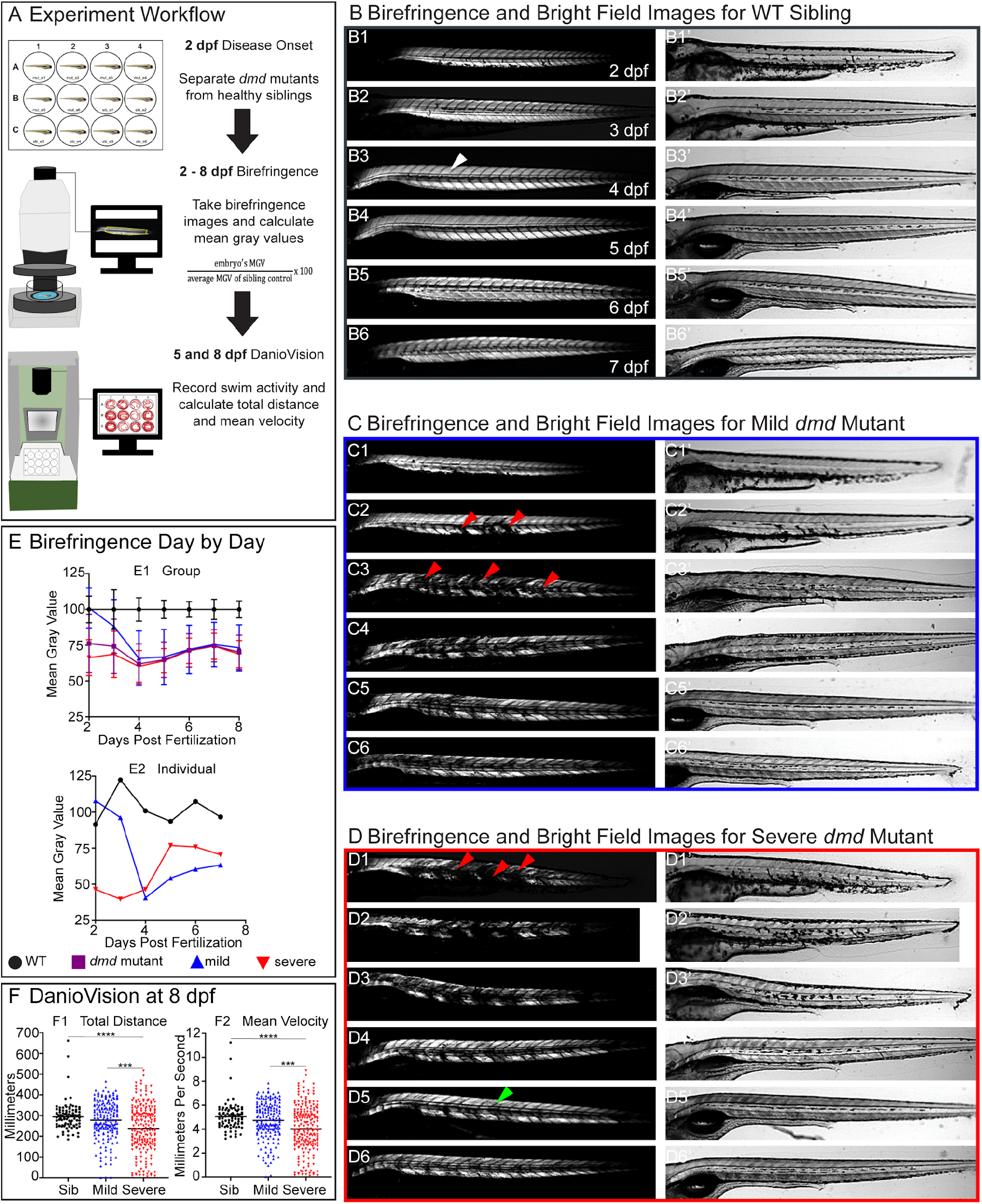
Variation in the *dmd* mutant phenotype determines disease progression. (A) We created an experiment workflow to assess disease progression from 2 to 8 dpf. At 2 dpf, birefringence is used to separate *dmd* mutants from WT siblings. Zebrafish are placed in individual wells of a 12-well plate and assigned a number, which is used to track individual zebrafish for the duration of the experiment. Each day, from 2 dpf to 8 dpf, birefringence images are taken. Birefringence (white) reflects normally organized muscle tissue. Loss of birefringence (grey to black) reflects areas of degeneration and myotomes with detached muscle fibers. Mean gray value is used to quantify birefringence and is presented as a percentage of WT sibling controls. Following birefringence imaging at 8 dpf, swimming activity is recorded using DanioVision. Total distance and mean velocity is calculated during the active (dark) periods. (B - D) Anterior left, dorsal top, side mounted. (B) Birefringence and bright field images of a WT sibling from 2 to 7 dpf. (C - D) At disease onset, zebrafish exhibit two levels of severity. (C) Birefringence and bright field images for a mild *dmd* mutant. Mild *dmd* mutants have mean gray values greater than 86% of WT siblings at 2 dpf. (D) Birefringence and bright field images for a severe *dmd* mutant. Severe *dmd* mutants have mean gray values less than 85.99% of WT siblings at 2 dpf. (E) Mild and severe *dmd* mutants exhibit variation in disease progression. (E1) Average mean gray values for siblings (black circles) do not change across time, remaining at 100%. However, mild *dmd* mutants (blue upward facing triangles) undergo extensive degeneration for the first three days followed by a period of slight regeneration. Conversely, severe *dmd* mutants (red downward facing triangles) regenerate throughout the study. (E2) Individual mean gray values for zebrafish presented in B - D highlight the vast degeneration that mild *dmd* mutants experience compared to the regeneration that occurs in severe *dmd* mutants. (F) Swimming activity is significantly different in mild versus severe *dmd* mutants. Total distance (F1) and mean velocity (F2) are significantly lower in severe *dmd* mutants compared to mild *dmd* mutants and WT siblings at 8 dpf. Each data point represents a single time point for an individual zebrafish. Each zebrafish has a total of 15 points. DanioVision data were analyzed using an ordinary one-way ANOVA with Tukey’s multiple comparisons test. *** p < 0.001, **** p < 0.0001.

### DanioVision Analysis

The DanioVision system and EthoVision XT 13.0 software (Noldus, Virginia, USA) was used to conduct high-throughput locomotion tracking studies to better characterize the impact of NMES on zebrafish swim function. A clear 12-well plate was placed into the DanioVision observation chamber. The temperature control unit was set to 28.5 degrees Celsius, ensuring that the temperature of the ERM in the well plate was maintained. Zebrafish had a 5 minute acclimation period to the observation chamber prior to the beginning of the recording period. Using the EthoVision software, we created a white-light routine that included 5 minutes in the dark followed by two light-on/off cycles, where the white light turned on at 100% intensity for 5 minutes and then turned off for 5 minutes. The total recording time was 25 minutes. Recordings were made at the same time each day. For each fish, the average total distance and mean velocity across 1 minute intervals were calculated, such that each fish had a total of 25 measurements for total distance and for mean velocity. We then focused on swim activity during the three 5-minute dark periods, which represent when zebrafish are most active. This analysis provided 15 measurements for each zebrafish.

### Membrane Permeability Indicated by Evan’s Blue Dye

Evan’s blue dye is a membrane impermeable dye used to assess membrane damage. In *dmd* mutants, EBD is used to assess muscle fiber integrity, and we used EBD to assess fiber integrity pre- and post-NMES using the methods described by (Smith et al., 2015). EBD (Sigma-Aldrich, Missouri, USA) was dissolved to 1% w/v in 0.9% saline solution. This EBD stock solution was further diluted to 0.1% then loaded into an injection needle pulled from glass capillary tubes on a Sutter Flaming/Brown Micropipette Puller (California, USA). Zebrafish were placed in tricaine (612 μM) for 4 minutes. At the end of the 4 minutes, zebrafish were aligned on a 1% agarose-lined Petri dish in a minimal volume of ERM. The needle was gently inserted into the peri-cardial space and EBD was injected using a MPPI-3 pressure injector (ASI, Eugene, Oregon). Zebrafish were allowed to recover for 4 hours, providing ample time for the dye to circulate the body and enter damaged muscle fibers. Zebrafish were prepared for live imaging as described above for birefringence. An ET DSR fluorescent filter (Leica) was used to visualize EBD. After imaging the initial dye amount in each zebrafish, zebrafish underwent 1 session of NMES as described above. Immediately after the NMES session, zebrafish were again prepared for live imaging. This allowed us to observe whether NMES caused additional dye entry into the muscle. Imaging parameters remained the same for all zebrafish and imaging sessions. Zebrafish were mounted laterally with the head on the left and dorsal up. To quantify EBD entry, we calculated mean gray values using the same methods described for birefringence except the outline was drawn from the first visible somite to the last visible somite. All images were blinded prior to analysis using a Perl script. Data is presented as the average mean gray value for three separate measurements.

### Immunostaining

Zebrafish were fixed in 4% paraformaldehyde for 4 hours at room temperature. After fixation, embryos were rinsed in PBS-0.1% Tween 20 (PBS-tw; BIO-RAD, Hercules, California). For visualizing muscle structure, phalloidin was used. Zebrafish were first permeabilized in PBS-2% Triton-X-100 (Fisher Scientific, Waltham, Massachusetts) for 1.5 hours and then placed in 1:20 phalloidin (Invitrogen, Eugene Oregon) in PBS-tw for 4 hours on the rocker at room temperature. Zebrafish were rinsed out of phalloidin using PBS-tw and stored in PBS-tw until imaged. For visualizing neuromuscular junctions, zebrafish were stained with alpha-bungarotoxin and SV2. Zebrafish were first permeabilized in 1 mg/ml collagenase in 1X PBS for 1.5 hours, and then stained with 1:500 alpha-bungarotoxin-647 (Invitrogen) and 1:20 phalloidin in antibody block (5% BSA (Fisher Scientific), 1% DMSO (Sigma-Aldrich), 1% Triton-X-100, 0.2% saponin from quillaja bark (Sigma-Aldrich) in 1X PBS) for 2 hours at room temperature. Zebrafish were rinsed using PBS-tw, and placed in antibody block overnight at 4 degrees Celsius. Zebrafish were then stained with 1:50 SV2 (DSHB, Iowa City, Iowa) in antibody block for 3 days at 4 degrees Celsius. Upon removal from SV2, zebrafish were rinsed using PBS-tw and then placed in antibody block for 8 hours on the rocker at room temperature. This was followed by an overnight incubation in 1:200 GAM (Invitrogen) in antibody block. Zebrafish were then rinsed out of secondary antibody using PBS-tw and stored in PBS-tw until imaged. Phalloidin-488 or −546 and GAM-488 or −546 were used interchangeably with no differences in staining observed.

### Imaging

Confocal imaging was used to visualize phalloidin and NMJ staining. Fixed and stained zebrafish were deyolked and then mounted in a 24-well glass bottom plate using 0.5% low-melt agarose (Boston BioProducts, Ashland, Massachusetts) in 1X PBS. For live confocal time-lapse imaging, zebrafish were anesthetized in tricaine solution (612 μM in 1X ERM) for 4 minutes and then mounted in a 24-well glass bottom plate or 30-mm glass bottom petri dish using 0.5% low-melt agarose in 1X ERM (with 612 μM tricaine). Two or three zebrafish were placed in each well. Zebrafish were mounted anterior left and dorsal up to ensure the same side of the fish was imaged each day. Finally, a small amount of tricaine solution (612 μM in 1X ERM) was added to prevent the agarose from evaporating and to ensure the zebrafish remained anesthetized throughout the imaging session. Upon completion of imaging, zebrafish were gently removed from the agarose using fine fishing line and returned to their respective wells. All fluorescent images were captured using the Leica SP8 confocal microscope.

Second harmonic generation (SHG) imaging was used as a label-free mechanism to visualize sarcomeres. Fixed zebrafish were deyolked and then mounted in a 30-mm glass bottom petri dish using 1.0% low-melt agarose in 1X PBS. Once the agarose solidified, the petri dish was filled with 1X PBS. Images were acquired using a custom-built two-photon microscope. This system uses a modified Olympus FV300 system with an upright BX50WI microscope stand and a mode-locked Ti:Sapphire (Coherent Ultra II) laser. Laser power was modulated via an electro-optic modulator. The SHG signals were collected in a non-descanned geometry using a single PMT. Emission wavelengths were separated from excitation wavelengths using a 665 nm dichroic beam splitter followed by a 448/20 nm bandpass filter for SHG signals. Images were acquired using circular polarization with excitation power ranging from 1 to 50 mW and a 40x 0.8 NA water immersion objective with 3x optical zoom and scanning speeds of 2.71 seconds per frame. All images were 512 × 512 pixels with a field of view of 85 micrometers.

### Image Analysis

All images were blinded using a Perl script prior to analysis. The percent of myotomes with muscle fiber detachments was calculated manually by counting the number of muscle segments with visibly detached fiber(s). Muscle segments are defined as half myotomes. Additionally, we used machine learning to identify healthy versus unhealthy muscle fibers. For these analyses, we used MATLAB to implement a deep learning approach to segment images of phalloidin stained fish into healthy muscle, sick muscle, and background. We used the DeepLab v3+ system with an underlying Resnet18 network (Chen et al., 2017). We defined the ground truth dataset manually using LabelBox (labelbox.com). Training images and ground truth images were broken down into 256 × 256 pixel images for training. The training dataset was divided into 60% training, 20% validation and 20% test data. Median frequency weighting was used to balance the classes. Each fish was oriented such that the head of the fish would be at the left of the image. Data was augmented to translate the images by 10 pixels vertically and horizontally. Rotation was found to make the network less accurate as orientation angle of the muscle fibers relative to the body orientation is important to assessing their health. The stochastic gradient descent with momentum (SGDM) optimizer was selected with 0.9 momentum. The maximum number of epochs was 100, and the mini-batch size was 8. In every epoch, the training dataset was shuffled. The number of iterations between evaluations of validation metrics was 315. The patience of validation stopping of network training is set up to 4. The initial learning rate used for training was 0.001. The learning rate was dropped 0.3 fold piecewise during training every 10 epochs. The Factor for L2 regularization (weight decay) was 0.005. The training set reached an accuracy of 97%. Images were then segmented by the MATLAB *semanticseg* command, which produced 8-bit unsigned integer segmentations. The fraction of each fish that was determined to be healthy was reported as a fraction of the total muscle. Pixels determined to be background (i.e. not muscle) were excluded from this calculation.

For NMJ analyses, we used a method previously published by our laboratory (Bailey et al., 2019). To prepare images for analysis, a custom Fiji macro was written to keep image processing consistent throughout all experiments. First, the raw .lif file was opened in FIJI and the image was split into its respective channels (phalloidin, AChR, and SV2). The phalloidin channel was immediately saved as a .tif file and closed. For the AChR and SV2 channels, duplicate z stacks were created and a 10-pixel radius Gaussian blur was applied. These blurred images were then subtracted from their original images, respectively. The resulting images were then merged to a single image and a maximum intensity projection was generated. This maximum intensity projection was saved as a .tif file and closed. For each experiment, the maximum intensity projections were combined into a single .tif file using a custom MATLAB script. This combined .tif file was then opened in FIJI and three separate masks, marking the fish body, horizontal myoseptum and myoseptal innervation, were drawn on the projected images using the Pencil tool. These masks were used to define individual muscle segments, where each muscle segment represents half of a single myotome. Using a custom MATLAB script, skeleton number and skeleton length were calculated for each muscle segment across all zebrafish analyzed.

Muscle nuclei were analyzed using FIJI’s 3D Objects Counter as well as the Measure tool. To prepare images for analysis we first reduced background noise by duplicating the z stack, performing a 10-pixel Gaussian blur on the duplicated image, and subtracting the blurred image from the original image. We then performed a 1-pixel Gaussian blur on the resultant image and set a threshold using ‘max entropy’ setting. With this image, we used the Analyze Particles tool to generate masks to use with the 3D Objects Counter tool as well as the Measure tool. The 3D Objects Counter tool provided surface area and volume measurements while the Measure tool provided perimeter, area, and major axis measurements, which were used to calculate filament index.

To calculate sarcomere lengths, SHG images were first imported into ImageJ, and then, using the Freehand selection tool, two lines were drawn to indicate the outer boundaries (top and bottom) of the muscle fiber being analyzed. The Freehand selections were converted into .txt files and imported into LabVIEW VI. Using LabVIEW, the midline of the two selections (top and bottom) was determined. The midline was then imported back into ImageJ over the original photo, such that it was positioned in the center of the sarcomeres. Next, the Plot Profile tool and Peak Finder tool were used to determine the peaks, which correspond to sarcomere length. Since the Peak Finder tool gives distance in pixels, a conversion factor was used to convert pixels to micrometers based on the objective and optical zoom used. Multiple muscle fibers are analyzed for each zebrafish. We avoided the optical illusion effect of ESH veneers regions when measuring sarcomere lengths (Dempsey et al., 2015).

### RNA Extraction and RNAseq

Total RNA was extracted from whole zebrafish at 7 dpf from replicate samples using the Zymo Direct-zol RNA microprep kit. Each biological replicate consisted of two zebrafish. For WT siblings, there were 4 replicates for the control group and 3 replicates for the eNMES group, and for *dmd* mutants, there were 8 replicates for the control group and 10 replicates for the eNMES group. Prior to performing RNA extractions, zebrafish within the eNMES and control groups were grouped based on their severity at disease onset and the calculated change in their birefringence from 5 dpf to 7 dpf. RNA was kept at −80 degrees Celsius until it was shipped to Quick Biology (Pasadena, California) for sequencing. Following RNA quality control using an Agilent BioAnalyzer 2100 (), polyA+ RNA-seq libraries were prepared for each sample using the KAPA Stranded RNA-Seq Kit (KAPA Biosystems, Wilmington, MA). Final library quality and quantity were analyzed by Agilent Bioanalyzer 2100 and Life Technologies Qubit3.0 Fluorometer. Each library was sequenced using 150 bp paired-end reads using an Illumina HiSeq4000 (Illumnia Inc., San Diego, CA).

Analyses of RNA-Seq reads were completed on the Advanced Computing Group Linux cluster at the University of Maine. To determine the quality of the RNA sequencing reads before further processing, FastQC version 0.11.7 was utilized. Following this quality assessment, reads were concatenated tail-to-head to produce one forward FASTQ file and one reverse FASTQ file for each sample. These FASTQ files were then trimmed of adapter sequences, and low quality leading and trailing ends were removed using Trimmomatic version 0.36.0 (Bolger et al., 2014). Trimmed paired-end reads mapped to the Ensembl-annotated zebrafish transcriptome (Ensembl version 95) to generate read counts per gene using RSEM version 1.2.31 (Li and Dewey, 2011) with bowtie version 1.1.2 (Langmead et al., 2009). Read counts were analyzed using the DESeq2 version 1.22.2 (Love et al., 2014) to analyze gene expression, p-value, and false discovery rate (FDR). Genes with fewer than ten mapped reads across all samples were excluded. For each pairwise comparison of treatment groups, differentially expressed genes were determined using FDR p-value cutoff of 0.1 and requiring at least a 0.6 log2 fold-change (in either direction). Resulting lists were used for Gene Ontology enrichment analysis and set analysis for each pairwise comparison.

Sets of differentially expressed genes (both increased and decreased expression) were analyzed to test for enriched GO Biological Process terms (FDR < 0.1) using GOrilla (http://cbl-gorilla.cs.technion.ac.il/). For this analysis, the entire set of expressed genes were used as a background. In cases where GOrilla found no enriched terms, PantherDB’s overrepresentation test on Biological Processes (http://pantherdb.org/) was used. Again, the entire set of expressed genes list was used as the background, and results were evaluated using Panther’s Fisher’s Exact Test and p-values were adjusted for multiple testing using FDR.

Ensembl gene IDs were mapped to gene symbols and names using zebrafishMine’s Analyse feature (http://www.zebrafishmine.org/). In some cases, manual mapping was used by comparing Zfin.org gene search and Ensembl gene search results. Summarized gene expression data are available at the Gene Expression Omnibus (accession number GSE155465), and FASTQ files are available at the Short Read Archive (accession number SRP274405).

### Cell Adhesion

Muscle fiber attachment strength was assessed similarly to that published by (Subramanian and Schilling, 2014). Zebrafish larvae were anesthetized with tricaine (612 μM in 1X ERM) for 4 minutes and then placed in the NMES gym. The stimulator settings were adjusted such that the frequency was 4 pulses per second, the delay was 60 ms, the duration was 2 ms, and the voltage was 30 volts. Zebrafish were stimulated for 1 minute. Birefringence images were taken pre- and post-stimulation as was described for the EBD study. Zebrafish were then subjected to a second round of stimulation, and birefringence images were taken after this second round (Figure 11B). Zebrafish were mounted laterally with the head on the left and dorsal up and the same imaging parameters were used for all zebrafish.

### Statistical Analysis

All statistical analyses were performed in Graphpad Prism. Normality was first assessed for all data using the Shapiro-Wilk test. If data passed this normality test, an unpaired two-tailed t test was performed between two data sets (i.e., *dmd* mutant control versus *dmd* mutant eNMES) while an ordinary one-way ANOVA was performed followed by a Tukey’s multiple comparison test between three data sets (i.e., WT sibling control versus *dmd* mutant control versus *dmd* mutant eNMES). Conversely, if data failed the Shapiro-Wilk normality test, a Mann-Whitney U test was performed for comparing two data sets while a Kruskal-Wallis test was performed for comparing three data sets. Significance for all tests was set to p < 0.05.

## RESULTS

### Longitudinal studies indicate that muscle structure, degeneration, and regeneration in *dmd* mutant zebrafish are variable

The zebrafish model of DMD has been used extensively in studies elucidating the underlying mechanisms of disease as well as screening for potential therapeutic compounds (Maves, 2014). We found that *dmd* mutant zebrafish exhibit some phenotypic variation. We used longitudinal birefringence imaging to track muscle structure and DanioVision to analyze swimming activity (as one metric of muscle function). We took two approaches to improve consistency in the data. For a given experiment, all birefringence imaging and DanioVision sessions were conducted at the same time every day. All birefringence data were also normalized to the average WT birefringence in each imaging session (Fig. 1A). Birefringence clearly visualizes healthy muscle in WT larvae (Fig. 1B, white arrowhead in B3 denotes a healthy muscle segment). While the average for a group of WT larvae is made 100% (see Fig. 1E1), birefringence of an individual WT larva varies but hovers around 100% Fig. 1D1). At the onset of muscle degeneration in *dmd* mutants, there was drastic variation in muscle structure; ranging from 35% to 135% of the average WT birefringence (Fig. 1E1, note the large standard deviation). Mutants were categorized as either mild or severe at the onset of muscle degeneration with mild having a mean gray value of ≥ 86% of WT birefringence and severe having a mean gray value of ≤ 85.999%. Although mild and severe mutants had indistinguishable muscle degeneration at 8 dpf (Fig. 1E1), they took different paths to get there. Mild *dmd* mutants had better muscle structure at 2 and 3 dpf than severe mutants (compare Fig. 1Cd, red arrowheads point to a couple muscle segments with degeneration versus the near complete degeneration in severe mutants in Fig. 1D1). This improved muscle structure was reflected in significantly higher mean gray values at 2 and 3 dpf compared to severe *dmd* mutants, but at 5 and 6 dpf, severe *dmd* mutants had significantly higher mean gray values. These data indicate that *dmd* mild mutants undergo extensive degeneration for the first three days after disease onset. Next, they undergo a period of slight regeneration. In contrast, muscle in severe *dmd* mutants regenerated throughout the study. These data clearly indicate that there is phenotypic variation in the zebrafish *dmd* mutant and that this variation can be quantified with birefringence.

As general muscle structure at 8 dpf was not significantly different in mild versus severe mutants we asked whether muscle function, as assayed by swimming activity, was different in mild versus severe mutants. Using DanioVision to capture swimming activity, we found that mild mutants swam a significantly greater distance (n = 12 mild, n = 14 severe; p = 0.0003) with significantly faster velocity (n = 12 mild, n = 14 severe; p = 0.0003) than severe mutants at 8 dpf (Figure 1F). This result indicates that muscle structure early in development (2 dpf) correlates with function much later in development (8 dpf).

### Extended inactivity improves swimming but decreases lifespan

Currently, individuals with DMD are advised to refrain from excessive activity beyond that of their daily living. The zebrafish model of *dmd* has fewer muscle detachments while anesthetized in tricaine (Berger et al., 2010), supporting the hypothesis that activity may progress the disease more quickly. However, only three studies have examined the long-term impact of inactivity on disease progression (Hourdé et al., 2013; Mizuno, 1992; Mokhtarian et al., 1999). Thus, we asked how intermittent versus extended inactivity affects *dmd* zebrafish muscle in a longitudinal study. The intermittent inactivity protocol included 12 hours in low-dose tricaine (a broad voltage-gated sodium channel blocker) for three consecutive nights beginning at the onset of muscle degeneration. Larvae were removed from tricaine for 12 hours during the day. After three nights of inactivity, larvae were reared for three more days. Birefringence was again used as a metric to assess changes in overall muscle structure from 5 to 8 dpf. A positive change in birefringence (mean gray value) meant that there was increased birefringence at 8 dpf compared to 5 dpf and thus muscle structure improved. Conversely, a negative change meant that there was decreased birefringence and that muscle structure deteriorated from 5 to 8 dpf. Three intermittent periods of inactivity did not affect muscle structure (Figure 2A1-6). Next, we fixed and stained larvae with phalloidin at 8 dpf to determine whether there were any dramatic changes in muscle fiber structure. This approach also indicated that intermittent inactivity did not have major effects on muscle fiber organization (Figure 2A7). Interestingly, however, larvae subjected to intermittent inactivity swam more slowly (n = 9 control, n = 9 inactive; p < 0.0001) and covered less distance (n = 9 control, n = 9 inactive; p < 0.0001) at 8 dpf (Figure 2A8 and A9). These data indicate that early intermittent inactivity can have negative impacts on swimming activity later in development. However, survival was not negatively impacted (Figure 2A10).

**Figure 2:**
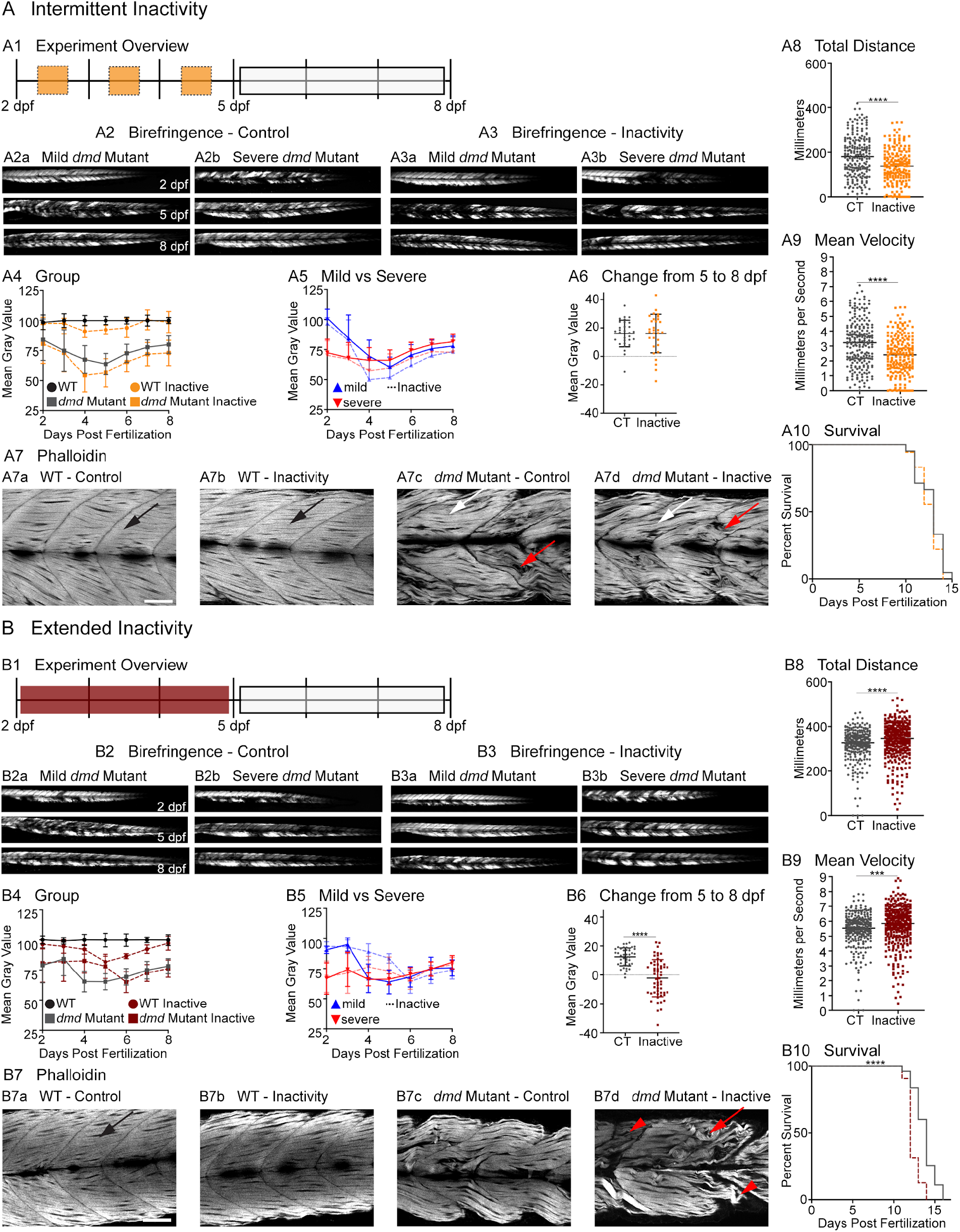
Inactivity in *dmd* mutants differentially affects muscle structure, function, and survival. (A1) Experiment overview for intermittent inactivity. Zebrafish are housed in a low dose of tricaine for 12 hours overnight (orange boxes) at 2, 3, and 4 dpf. Upon removal from tricaine at 5 dpf, zebrafish are allowed to recover in ERM (white box) for the remainder of the experiment. (A2 - A3) Anterior left, dorsal top, side mounted. (A2) Birefringence images at 2, 5, and 8 dpf for mild (A2a) and severe (A2b) *dmd* mutant controls housed in ERM. (A3) Birefringence images at 2, 5, and 8 dpf for mild (A3a) and severe (A3b) *dmd* mutants housed in tricaine for 12 hours overnight for 3 nights. (A4) Average mean gray values for WT sibling controls (black circles) remain consistent across time. However, WT siblings that were inactive (orange circles) experience a decrease in mean gray values at 4, 5 and 6 dpf, but recover to WT sibling control values by 8 dpf. Conversely, *dmd* mutants that were inactive (orange squares) experience a decrease in mean gray values compared to *dmd* mutant controls (gray squares) beginning at 4 dpf but do not recover to *dmd* control values at 8 dpf. (A5) Mild (blue upward facing triangles) and severe (red downward facing triangles) *dmd* mutants that were inactive (dashed lines) have lower mean gray values compared to their respective controls (solid lines) beginning at 4 dpf. Inactive mild *dmd* mutants experience a more dramatic decrease in mean gray values compared to mild control and inactive severe *dmd* mutants. (A6) Change in mean gray value from 5 to 8 dpf is not different in control versus inactive *dmd* mutants. (A7) Anterior left, dorsal top, side mounted. Scale bar is 50 micrometers. Phalloidin staining at 8 dpf suggests no change in muscle fiber structure following inactivity in *dmd* mutants (A7d) compared to *dmd* mutant controls (A7c). Total distance (A8) and mean velocity (A9) at 8 dpf are significantly lower in inactive *dmd* mutants compared to *dmd* mutant controls. Each data point represents a single time point for an individual zebrafish. Each zebrafish has a total of 15 points. (A10) Survival is not affected by intermittent inactivity. (B1) Experiment overview for extended inactivity. Zebrafish are housed in a low dose of tricaine (dark red box) for 72 hours beginning at 2 dpf. Upon removal from tricaine at 5 dpf, zebrafish recover in ERM (white box) for the remainder of the experiment. (B2) Birefringence images at 2, 5, and 8 dpf for mild (B2a) and severe (B2b) *dmd* mutant controls housed in ERM. (B3) Birefringence images at 2, 5, and 8 dpf for mild (B3a) and severe (B3b) *dmd* mutants housed in tricaine for 72 hours. (B4) Average mean gray values for WT sibling controls (black circles) remain consistent across time at 100%. However, WT siblings that were inactive (dark red circles) experience a dramatic decrease in mean gray value beginning at 4 dpf and continuing through 7 dpf, but return to control values by 8 dpf. Conversely, *dmd* mutants that were inactive (dark red squares) have higher mean gray values at 4 and 5 dpf compared to *dmd* mutant controls (gray squares). However, upon return to ERM, inactive *dmd* mutants experience a decrease in mean gray values but regenerate to *dmd* mutant control values. (B5) Mild (blue upward facing triangles) and severe (red downward facing triangles) *dmd* mutants that were inactive (dashed lines) have higher mean gray values compared to control *dmd* mutants (solid lines) at 4 and 5 dpf. Inactive mild *dmd* mutants experience a more dramatic decrease in mean gray values compared to inactive severe *dmd* mutants at 6 dpf. (B6) Change in mean gray value from 5 to 8 dpf is significantly lower in inactive versus control *dmd* mutants. (B7) Anterior left, dorsal top, side mounted. Scale bar is 50 micrometers. Phalloidin staining at 8 dpf suggests that inactivity negatively affects muscle structure in *dmd* mutants (B7d) compared to *dmd* mutant controls (B7c). Total distance (B8) and mean velocity (B9) at 8 dpf are significantly higher in inactive *dmd* mutants compared to *dmd* mutant controls. Each data point represents a single time point for an individual zebrafish. Each zebrafish has a total of 15 points. (B10) Survival is negatively affected by extended inactivity. Birefringence and DanioVision data were analyzed using two-sided t tests. Survival data were analyzed using a Mantel-Cox test. *** p < 0.001, **** p < 0.0001.

In contrast to intermittent inactivity, extended inactivity from 2 dpf through the morning of 5 dpf had deleterious impacts later in development. Immobilized larvae initially show improved muscle structure at 5 dpf (compare Fig. 2B3a to Fig. 2B2a, and Fig. 2B3b to Fig. 2B2b, quantification in Fig. 2B4). However, this improvement was short lived: there was a significant decline in birefringence of immobilized larvae 3 days after removal from tricaine (compare Fig. 2B3 5 dpf to 8 dpf, quantified in Fig. 2B4). These data indicate that while muscle structure was preserved during the inactive period, it became more susceptible to damage upon reinstatement of activity, which is indicated by the significant decrease (n = 43 control, n = 55 inactive; p < 0.0001) in mean gray value from 5 to 8 dpf (Fig. 2B6) and poor muscle fiber organization (Fig. 2B7d, red arrowheads denote short disorganized muscle segments, red arrow points to a degenerating fiber). Further, immobilized larvae swam a significantly lower total distance and at a significantly lower mean velocity compared to control *dmd* zebrafish 4 hours after removal from tricaine at 5 dpf (not shown). After three days of recovery in ERM at 8 dpf, swim function in inactive *dmd* zebrafish was significantly improved with these larvae swimming a significantly higher total distance (n = 19 control, n = 24 inactive; p < 0.0001) and at a significantly higher mean velocity (n = 19 control, n = 24 inactive; p < 0.0001) (Figure 2B8 and B9). Strikingly, this improved swimming activity did not correlate with survival: although extended inactivity increased swimming at 8 dpf, survival was negatively impacted (Figure 2B10). Thus, despite overall neutral effects on muscle structure at 8 dpf and improved swimming at 8 dpf, extended inactivity decreases lifespan (n = 55 control, n = 32 inactive; p < 0.0001).

### Extended inactivity diminishes *dmd* muscle resilience to neuromuscular electrical stimulation

The above data confirm previous data that inactivity improves muscle structure in *dmd* mutants while they remain inactive. However, our data show that this beneficial effect does not perdure. These results raise the question of why the seemingly improved muscle structure is not stable. To answer this question we turned to neuromuscular electrical stimulation (NMES), which was previously adapted for use in zebrafish larvae (Subramanian and Schilling, 2014). NMES uses trains of electrical pulses to evoke muscle contractions (Sheffler and Chae, 2007) and thus allows comparison of muscle structure in multiple larvae subjected to the same stimulus. Specifically, NMES would allow us to determine whether extended inactivity (1) obscured latent defects in muscle resilience because the muscle was not being used and thus did not degenerate, or (2) initially improved muscle fiber resilience but this resilience was not permanent. In order to distinguish between these possibilities, we asked whether inactive larvae were uniquely susceptible to activity (NMES) immediately upon removal from tricaine. We predicted that if inactivity obscured latent defects in muscle resilience, then NMES immediately after removal from tricaine would cause dramatic muscle damage. In contrast, if inactivity initially improved muscle fiber resilience, then NMES immediately after removal from tricaine would not cause muscle damage. For this experiment, larvae were placed in tricaine for three days and a birefringence image was taken immediately prior to removal from tricaine. This image served as the baseline birefringence level (see Fig. 3A). Next, larvae were subjected to a session of NMES. We used two different NMES paradigms, the first defined by high frequency/low voltage (NMES Paradigm 1), and the second defined by low frequency/high voltage (NMES Paradigm 2). Birefringence was imaged after this NMES session. Finally, larvae resumed normal activity for three days of recovery similar to the above experiments. At the end of the recovery period (8 dpf) we took birefringence images. Control *dmd* mutants, which were removed from tricaine and allowed to swim while experimental larvae were receiving NMES, sometimes showed increased degeneration just after swimming (Fig. 3B2b after) compared to immediately prior to removal from tricaine (Fig. 3B2a before). However, at a population level although there was a slight trend that inactive larvae showed more damage, there was not a significant increase in degeneration (Fig. 3B3). In contrast, both NMES paradigms significantly worsened mean gray values (NMES Paradigm 1: n = 8 ERM, n = 11 inactive, p = 0.0227; NMES Paradigm 2: n = 8 ERM, n = 12 inactive, p = 0.0157) immediately after stimulation in larvae that underwent three days of inactivity (Fig. 3C2, D2, yellow asterisks denote new areas of degeneration after NMES, Fig. 3C3 and D3). Larvae that underwent three days of inactivity exhibited more fiber detachments, especially following NMES Paradigm 2 (Fig. 3F,G, red arrows denote fiber detachments). These data suggest that inactivity for an extended period of time may cause *dmd* muscle to become more susceptible to contraction-induced injury.

**Figure 3:**
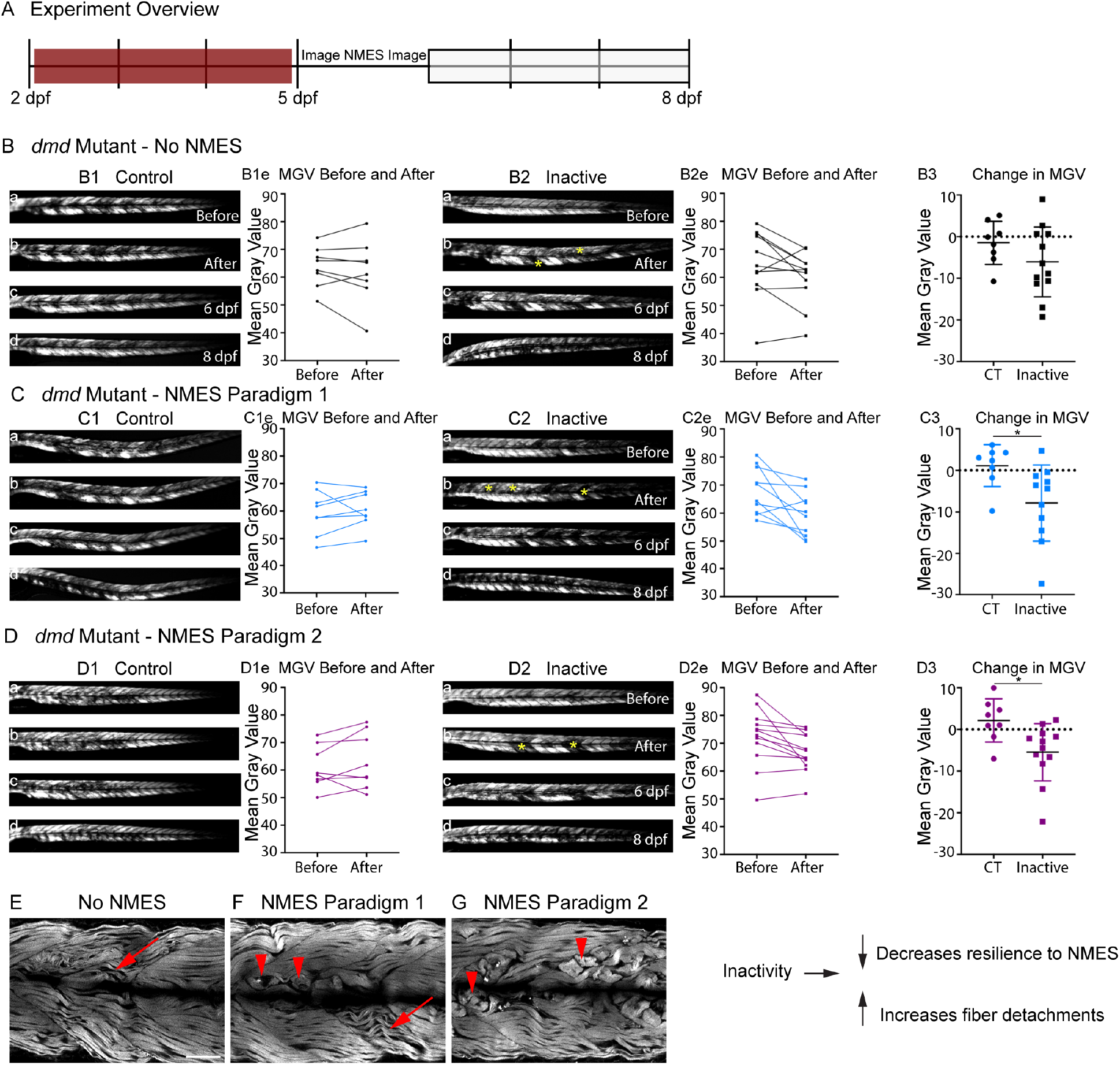
Extended inactivity increases susceptibility to injury in *dmd* mutants. (A) Experiment overview. Zebrafish are reared in a low dose of tricaine for 72 hours (dark red box) beginning at 2 dpf. At 5 dpf, zebrafish receive a single session of NMES (either Paradigm 1 or Paradigm 2) and are then allowed to recover in ERM (white box) for the remainder of the experiment. (B, C, D) Anterior left, dorsal top, side mounted birefringence images. Yellow asterisks indicate degeneration in the “after” panels. (B1) Birefringence for the first (a) and second (b) imaging session, and at 6 (c) and 8 (d) dpf for a *dmd* mutant control that did not receive NMES. (B1e) Individual mean gray values for *dmd* mutant controls for the first and second imaging session. (B2) Birefringence for the first (a) and second (b) imaging session, and at 6 (c) and 8 (d) dpf for an inactive *dmd* mutant that did not receive NMES. (B2e) Individual mean gray values for inactive *dmd* mutants for the first and second imaging sessions. (B3) Change in mean gray values between the first and second imaging session are lower in the inactive *dmd* mutants, indicating that upon removal from tricaine, muscle immediately begins degenerating. (C1) Birefringence before (a) and after (b) NMES Paradigm 1, and at 6 (c) and 8 dpf for a *dmd* mutant control. (C1e) Individual mean gray values for *dmd* mutant controls before and after NMES Paradigm 1. (C2) Birefringence before (a) and after (b) NMES Paradigm 1, and at 6 (c) and 8 (d) dpf for an inactive *dmd* mutant. (C2e) Individual mean gray values for inactive *dmd* mutants before and after NMES Paradigm 1. (C3) Change in mean gray values before versus after NMES Paradigm 1 are significantly lower in inactive *dmd* mutants, indicating *dmd* muscle fibers are less resilient following extended inactivity. (D1) Birefringence before (a) and after (b) NMES Paradigm 2, and at 6 (c) and 8 (d) dpf for a *dmd* mutant control. (D1e) Individual mean gray values for *dmd* mutant controls before and after stimulation. (D2) Birefringence before (a) and after (b) NMES Paradigm 2, and at 6 (c) and 8 (d) dpf for an inactive *dmd* mutant. (D2e) Individual mean gray values for inactive *dmd* mutants before and after NMES Paradigm 2. (D3) Change in mean gray values before versus after NMES Paradigm 2 are significantly lower in inactive *dmd* mutants, further indicating that muscle fibers are less resilient after inactivity. (E-G) Anterior left, dorsal top, side mounted phalloidin staining. Red arrows denote disorganized fibers, red arrowheads point to detached fibers. A single session of NMES Paradigm 1 negatively affects muscle structure in inactive *dmd* mutants (F) compared to inactive *dmd* mutants that did not receive stimulation (E). Similarly, a single session of NMES Paradigm 2 negatively affects muscle structure in inactive *dmd* mutants by increasing the number of visibly detached fibers (G) compared to inactive *dmd* mutant controls. Each data point represents a single zebrafish. Birefringence data were analyzed using two-sided t tests. * p < 0.05, ** p < 0.01. Scale bar is 50 micrometers.

### A model for studying the impact of NMES on *dmd* muscle

Strength training is an excellent approach to combat muscle wasting and weakness in healthy individuals. Using zebrafish larvae as a model for lifting weights is not feasible, so we asked whether we could use NMES as an alternate means to stimulate muscle activity and combat muscle wasting and weakness in *dmd* larvae. Previous work used NMES as a controlled stimulus designed to illuminate the fact that muscle that appeared morphologically normal was actually highly susceptible to stimulation-induced injury (Subramanian and Schilling, 2014). We approached NMES from the opposite perspective and asked whether different NMES paradigms would have different impacts on neuromuscular stability in *dmd* larvae. The parameters that can be adjusted in NMES include pulse frequency and voltage. We designed four different NMES paradigms ranging from high frequency/low voltage pulse trains to lower frequency/higher voltage pulse trains (Figure 4C and D). In order to easily differentiate these paradigms from each other, and because they were conceptually based on strength training paradigms that vary in the number of repetitions and load, we named these paradigms endurance-NMES (eNMES), hypertrophy-NMES (hNMES), strength-NMES (sNMES), and power-NMES (pNMES). We first asked whether these different NMES paradigms elicited unique tail bend patterns that vary in how many times the tail bends as well as how hard it bends. As would be expected, eNMES with high frequency/low voltage pulse trains elicited a fast but subtle tail beat. Conversely, with pNMES, the tail beat infrequently but bent to a much greater degree (not shown).

**Figure 4:**
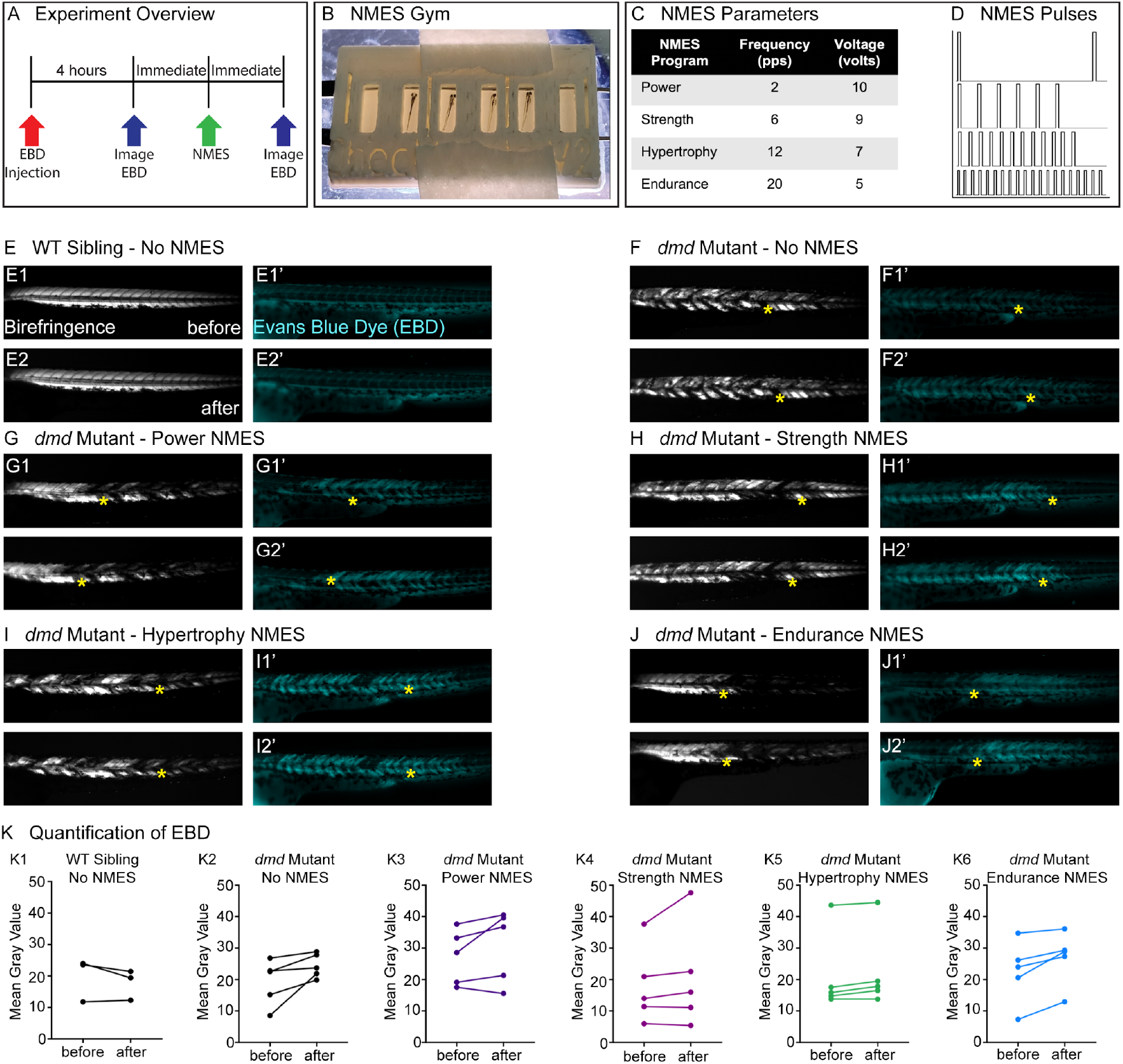
Four NMES paradigms do not result in immediate damage to the sarcolemma. Experiment overview. At 2 dpf, WT siblings and *dmd* mutants were injected with Evans blue dye (EBD). Four hours later, zebrafish were imaged for birefringence and EBD before and after a single session of NMES. (B) For NMES, zebrafish are placed in a 3D printed gym with their heads towards the positive electrode and tails towards the negative electrode. (C - D) NMES delivers a series of square wave pulses that vary in frequency and voltage. We named these paradigms after weightlifting regimes. (E-J) Anterior left, dorsal top, side mounted birefringence and EBD fluorescent images. Yellow asterisks denote the same position in embryos before and after NMES. (E) WT sibling control exhibits healthy muscle segments (E1, E2) and no dye entry in the muscle (E1’, E2’) during the first and second imaging session. (F) *dmd* mutant control has significant areas of degenerated muscle (F1) and dye entry (F1’) but no new areas of degeneration or dye entry during the second imaging session (F2, F2’). (G-J) Similar to the *dmd* mutant control, *dmd* mutants that receive NMES have significant areas of degenerated muscle and dye entry prior to NMES but no new areas of degeneration or dye entry during following NMES. (K) Quantification of EBD during the first and second imaging session.

### NMES does not result in immediate damage to the sarcolemma

One of the major reasons why strength training is not recommended for individuals with DMD is due to the fragility of the sarcolemma and its susceptibility to contraction-induced damage. Prior to evaluating each NMES paradigm, we asked whether the pulse parameters resulted in dramatic immediate damage to the sarcolemma. We did this by asking whether increased Evans blue dye (EBD) was observed in muscle after one session of NMES. EBD was injected into the peri-cardial space at 2 dpf and allowed to circulate for 4 hours. Then, images of EBD in the zebrafish trunk musculature were taken immediately prior to and after one session of NMES (Figure 4A). The relative amount of EBD in muscle was calculated using mean gray values of the EBD channel prior to and after NMES. Birefringence and EBD images of the same embryos before and after NMES are shown in Fig. 4. The yellow stars denote the same position in the embryo before and after stimulation. Both WT and *dmd* mutant control larvae are similar when imaged prior to and after the experimental larvae received NMES (Fig. 4E,F,K1,K2). None of the NMES paradigms consistently caused a dramatic change in either birefringence (not shown) or EBD infiltration (Fig. 4G-J, K, n = 5 embryos imaged, subjected to NMES, and imaged). These results indicate that the four NMES paradigms do not cause immediate dramatic damage to the sarcolemma.

### Different NMES paradigms differentially impact *dmd* muscle structure, function and survival

After determining that NMES paradigms did not cause immediate dramatic damage, we asked whether different NMES programs had different effects on the progression of muscle degeneration in *dmd* larvae. We developed a protocol that was divided into two periods: the training period and the recovery period (Fig. 5A). During the training period zebrafish completed three sessions of NMES, each session lasting one minute, on three consecutive days (2, 3, and 4 dpf) at the same time each day. Following these three training days, zebrafish entered the recovery period (5, 6, 7 and 8 dpf). Again, we evaluated muscle structure using birefringence and swim function using DanioVision. Therefore, the only aspects that changed across experiments were the NMES pulse parameters.

**Figure 5:**
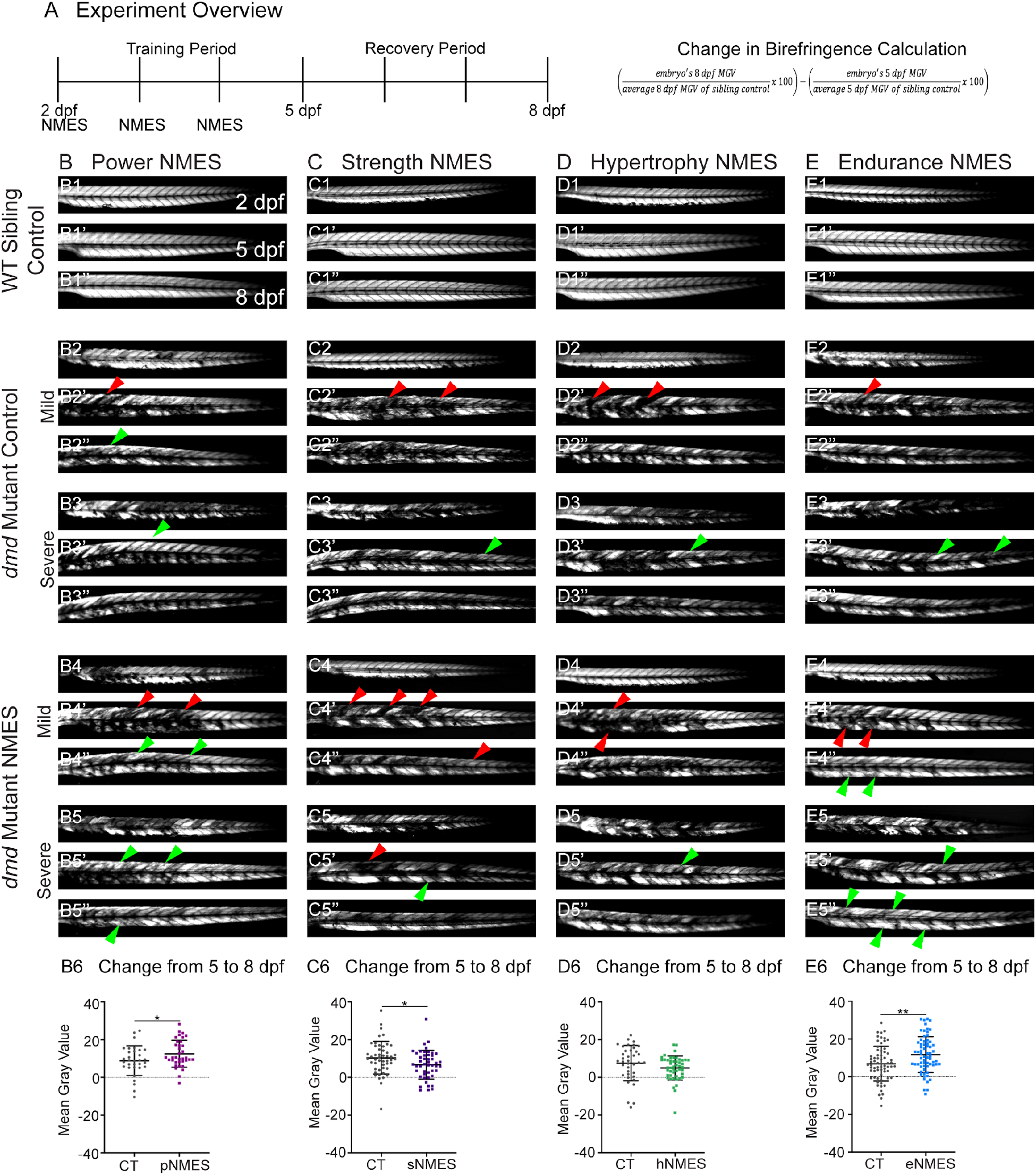
Birefringence is used as an initial measure of muscle structure following NMES. (A) Experiment overview and calculation of change in mean gray value from 5 to 8 dpf. At 2 dpf, birefringence images are taken followed by the first session of NMES. At 3 and 4 dpf, zebrafish undergo the second and third sessions of NMES, respectively. Birefringence images are taken at 5 and 8 dpf. The training program is divided into the training period (2 to 4 dpf) and the recovery period (5 to 8 dpf). (B - E) Anterior left, dorsal top, side mounted birefringence images for WT sibling controls (B1 - E1), mild (B2 - E2) and severe (B3 - E3) *dmd* mutant controls, and mild and severe *dmd* mutants that received pNMES (B4 and B5), sNMES (C4 and C5), hNMES (D4 and D5), or eNMES (E4 and E5). (B6, C6, D6, and E6) Change in mean gray values from 5 dpf to 8 dpf represent how the muscle responds to and recovers from 3 sessions of NMES. Positive changes indicate improvements in muscle structure while negative changes indicate deterioration in muscle structure. Red arrowheads denote degeneration from the previous time point, green arrowheads denote regeneration from the previous time point. Power (B6, maroon squares) and endurance (E6, blue squares) NMES significantly improve muscle structure in *dmd* mutants compared to *dmd* mutant controls (gray circles). Strength (C6, purple squares) NMES significantly worsens muscle structure in *dmd* mutants while hypertrophy NMES (D6, green squares) trends to decrease muscle structure compared to *dmd* mutant controls (gray circles). Each data point represents a single zebrafish. Birefringence data were analyzed using two-sided t tests. * < 0.05, ** p < 0.01.

As shown above for inactivity, we focused on the change in mean gray value from 5 dpf to 8 dpf because that change represents how the muscle responds to and recovers from 3 sessions of NMES. Wild-type larvae with all 4 NMES paradigms were unaffected (Fig. 5B1,C1,D1,E1, and data not shown). Control *dmd* larvae for each NMES paradigm were similar to larvae shown in Fig. 1, with mild larvae degenerating between days 2-5 (Fig. 5B2,C2,D2,E2, red arrowheads denote degeneration from the previous time point, green arrowheads denote regeneration from the previous time point); and severe larvae regenerating between days 2-5 (Fig. 5 B3,C3,D3,E3). Between days 5-8, as shown in Fig. 1, birefringence levels for both mild and severe larvae trend towards slight improvement (Fig. 5B6, C6, D6, E6). The eNMES and pNMES paradigms improved muscle structure in *dmd* mutants. pNMES resulted in a slight but significant increase in birefringence compared to controls (Fig. 5B6, note also green arrows in Fig. 5B4 and 5B5). eNMES also increased regeneration between 5 and 8 dpf (n = 66 control, n = 66 eNMES; p = 0.0037) (Fig. 5E6, note green arrows in Fig. 5E4, E5). In contrast, *dmd* mutants that underwent sNMES exhibited significantly lower changes in mean gray values compared to control *dmd* mutants (n = 49 control, n = 47 sNMES; p = 0.0302) (Figure 5C6) while hNMES trended towards lowering birefringence (Figure 5D6). These data indicate that, at least at a gross level, different NMES paradigms do have different effects on muscle structure in zebrafish larvae.

Muscle fibers in WT zebrafish are highly organized and linear (Figure 6A). In contrast, many fibers in *dmd* mutants are disorganized while others are compressed and/or detached from their extracellular matrix (Figure 6B). We quantified the percentage of muscle segments with fiber degeneration and found that, similar to the results observed with birefringence, eNMES (n = 22 control, n = 18 eNMES; p = 0.0191) and pNMES (n = 12 control, n = 20 pNMES; p = 0.0033) resulted in fewer fibers degenerating compared to control *dmd* mutants (Figure 6C2 and F2). Taken together, the above data indicate that eNMES and pNMES improve muscle structure in *dmd* larvae.

**Figure 6:**
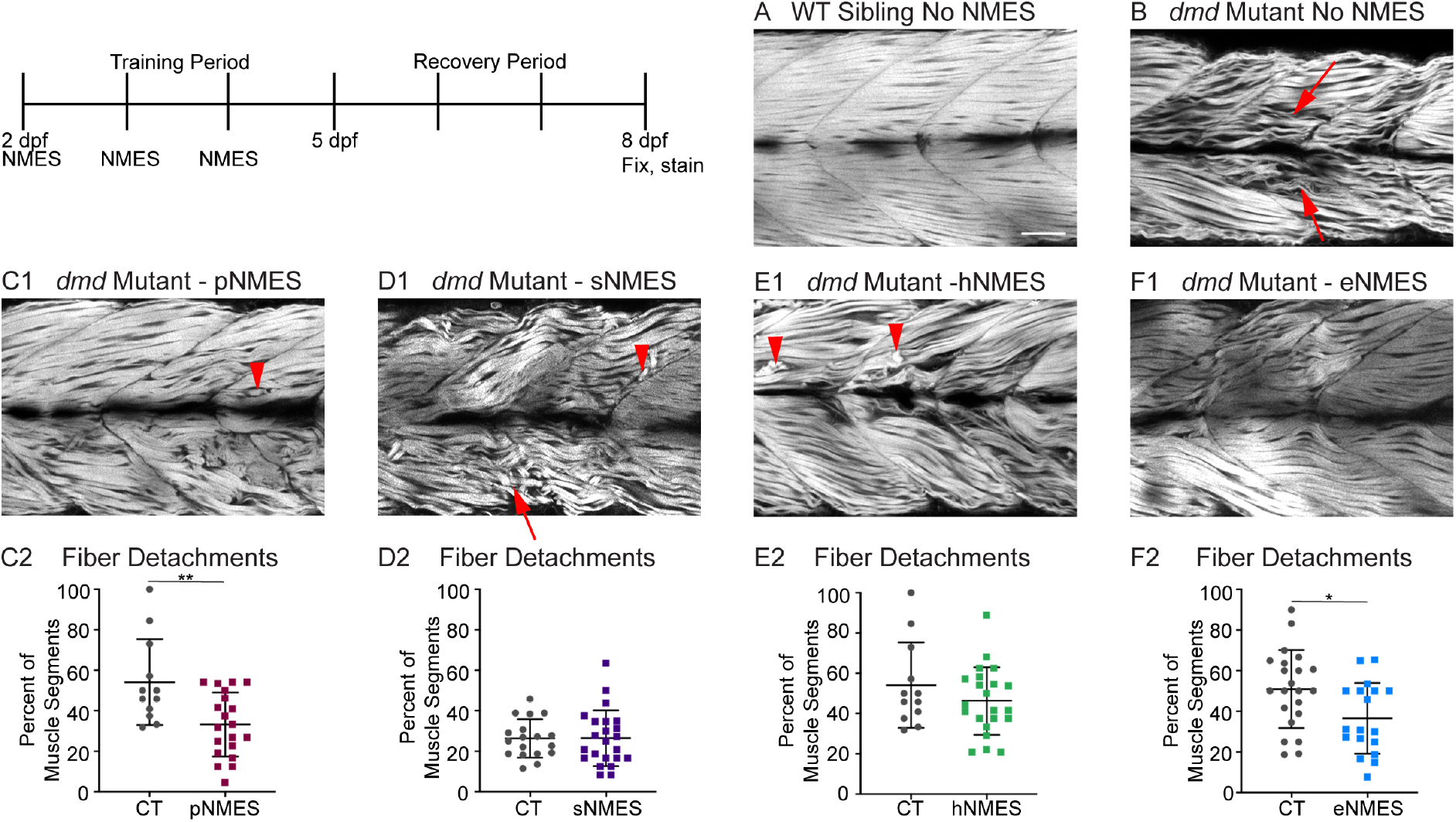
Phalloidin staining provides more details on how *dmd* muscle responds to NMES at the structural level. Phalloidin staining for F-actin at 8 dpf allows for visualization of individual muscle fibers and the ability to count detached fibers in *dmd* mutants. Anterior left, dorsal top, side mounted. Red arrows point to disorganized muscle fibers, red arrowheads point to detached muscle fibers. (A) Representative image of WT sibling demonstrates organized muscle fibers with well-defined myotome boundaries. (B) Representative image of *dmd* mutants demonstrates disorganized, wavy muscle fibers with poorly defined myotome boundaries and empty space between individual muscle fibers. (C1) Representative image of *dmd* mutant that received pNMES demonstrates less muscle fiber waviness, lack of empty space between muscle fibers but visible detached fibers. (D1) Representative image of *dmd* mutant that received sNMES demonstrates massive deterioration of muscle fiber structure, disorganized myotomes with poorly defined boundaries. (E1) Representative image of *dmd* mutant that received hNMES demonstrates improved muscle fiber organization with more defined myotome boundaries but visibly detached muscle fibers and empty space between fibers. (F1) Representative image of *dmd* mutant that received eNMES demonstrates healthy myotomes with clearly defined boundaries, organized muscle fibers with very few wavy fibers, and lack of empty space between fibers. Quantification of the percentage of muscle segments with detachments indicates that pNMES (C2) and eNMES (F2) significantly reduce fiber detachments in *dmd* mutants. Strength NMES (D2) and hNMES (E2) do not impact the percent of muscle segments with detachments. Each data point represents a single fish. A muscle segment is defined as half of a myotome. Muscle detachment data were analyzed using two-sided t tests.* p < 0.05, ** p < 0.01. Scale bar is 50 micrometers.

We hypothesized that improved muscle structure would correlate with improved function. We tested this hypothesis by assessing swim activity as a gross readout of muscle function. Swim activity was tested using DanioVision. Swimming activity was recorded for 25 minutes with alternating 5 minute light/dark periods. As predicted, eNMES resulted in increased distance (n = 7 control, n = 11 eNMES; p < 0.0001) and mean velocity (n = 7 control, n = 11 eNMES; p < 0.0001) compared to control *dmd* larvae (Figure 7A4 and B4). Surprisingly, though, pNMES negatively affected swimming activity (for both total distance and mean velocity: n = 20 control, n = 19 pNMES; p < 0.0001) (Figure 7A1 and B1) despite having improved muscle structure. sNMES also significantly reduced total distance (n = 26 control, n = 23 sNMES; p < 0.0001) and mean velocity (n = 26 control, n = 23 sNMES; p < 0.0001) (Figure 7A2 and B2) while hNMES did not affect these two measures (Figure 7A3 and B3).

**Figure 7:**
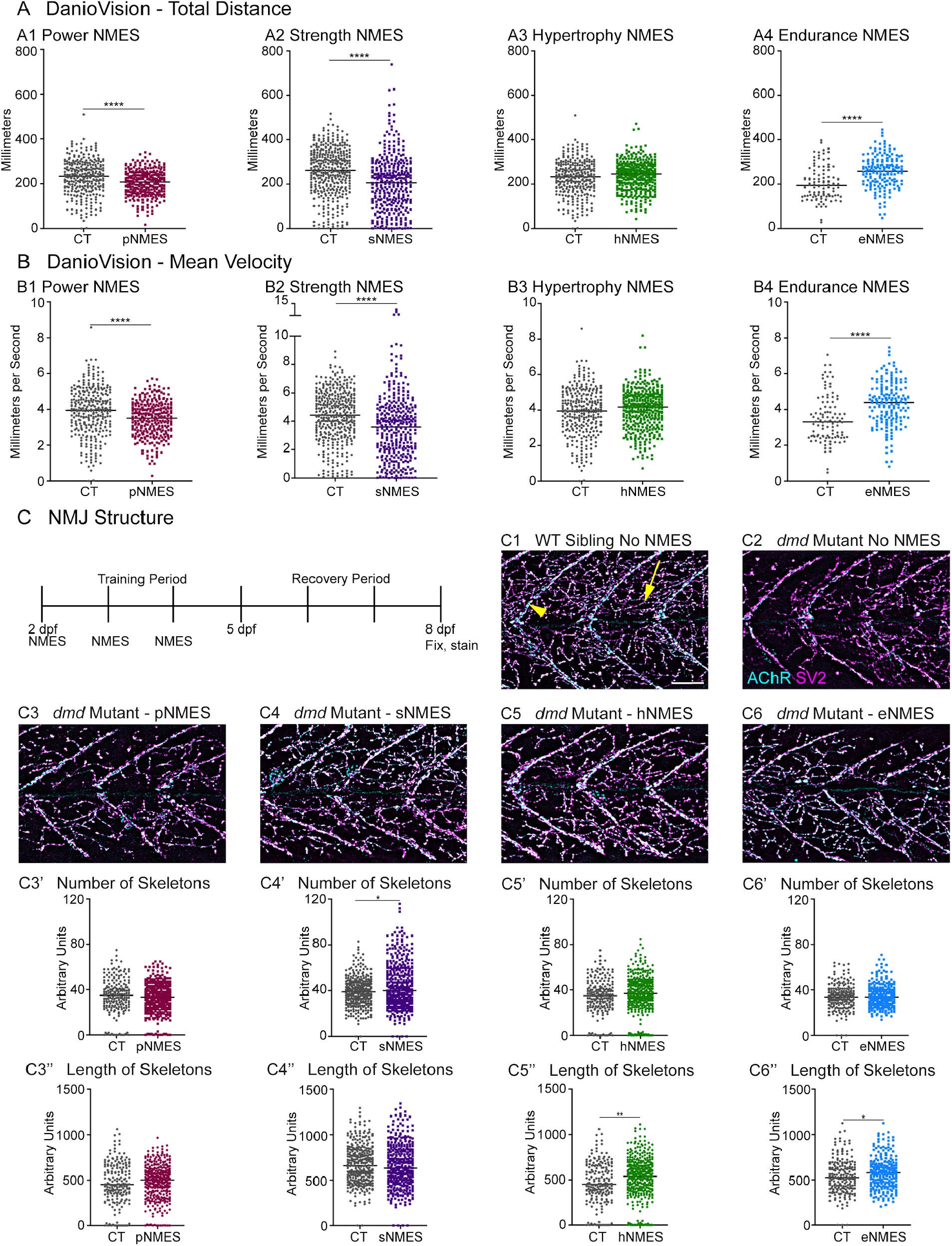
NMJ abundance does not correlate with swim function. DanioVision was used to assess the impact of NMES on total distance (A) and (B) mean velocity. Measurements were made at 8 dpf. (A1, B1) *dmd* mutants that completed pNMES exhibited significant reductions in total distance and mean velocity compared to *dmd* mutants in the control group. (A2, B2) Strength NMES also negatively affected swimming activity in *dmd* mutants compared to control *dmd* mutants. (A3, B3) No change in total distance or mean velocity is observed following hNMES. (A4, B4) *dmd* mutants that completed eNMES swam a significantly greater total distance and at a significantly faster mean velocity compared to *dmd* mutants in the control group. Each data point represents a single time point for an individual zebrafish. Each zebrafish has a total of 15 points. (C) anti-SV2 (cyan) a-Bungarotoxin (AChR; magenta) visualize the pre- and post-synaptic components of the NMJ, respectively. (C1) Representative image of WT sibling. Myoseptal innervation, innervation at the chevron-shaped myotendinous juncion (yellow arrowhead) is slow-twitch innervation. Fast-twitch muscle innervation is the network between the MTJs, yellow arrow points to fast-twitch muscle innervation. (C2) Representative image of *dmd* mutant demonstrates a visible reduction in innervation, with relatively large portions of the muscle segments lacking innervation. (C3, C4, C5, and C6) Representative images of *dmd* mutants that completed three sessions of sNMES, hNMES, or eNMES demonstrate increased innervation. NMJ images were skeletonized as previously described. sNMES increases the number of skeletons (C4’), whereas hNMES and eNMES increase skeleton length (C5’, C6’). Power NMES did not change the number or length of skeletons compared to *dmd* mutant controls. DanioVision data were analyzed using two-sided t tests. NMJ data were analyzed using either an ordinary one-way ANOVA with Tukey’s multiple comparisons test or a Kruskal-Wallis test with Dunn’s multiple-comparison test.** p < 0.01, *** p < 0.001, **** p < 0.0001.

Because improvements in muscle structure in response to different NMES paradigms did not strictly correlate to changes in swimming, we asked whether neuromuscular junction (NMJ) morphology changed with NMES. We analyzed NMJ morphology by using the SV2 antibody to label presynaptic structures and alpha-bungarotoxin to stain postsynaptic AChR. We focused on analyzing fast-twitch muscle fiber innervation, which is called distributed innervation (the rich network of NMJs, yellow arrow Fig. 7C1, in between the chevron shaped slow-twitch muscle innervation at the myotendinous junctions (MTJs, yellow arrowhead, Fig. 7C1)). Analysis was done in a semi-automated fashion that involved skeletonizing the NMJs as previously described (Bailey et al., 2019). We found that sNMES increased the number of skeletons (n = 18 control, n = 20 sNMES; p = 0.0395). In contrast, both hNMES and eNMES did not increase the number of skeletons but increased skeleton length (n = 15 control, n = 23 hNMES, p = 0.0036; n = 18 control, n = 15 eNMES, p = 0.0278). Thus, all NMES paradigms other than pNMES improved number or length of NMJs compared to control *dmd* mutants.

Lastly, survival was tracked in *dmd* mutants treated with NMES. Survival checks were performed twice daily. Three sessions of eNMES (n = 32 control, n = 37 eNMES; p < 0.0001), sNMES (n = 63 control, n = 55 sNMES; p = 0.0004), and pNMES (n = 37 control, n = 32 pNMES; p = 0.0414) slightly but significantly extended the median age of survival for *dmd* mutants compared to unstimulated *dmd* mutants (Figure 8A, B, D); with eNMES having the largest beneficial effect. Three sessions of hNMES, however, did not affect median survival age (Figure 8C). Taken together, these data indicate that different NMES paradigms elicit different neuromuscular responses. Furthermore, out of the four NMES paradigms we tested, only eNMES improves neuromuscular structure, swimming, and lifespan.

**Figure 8:**
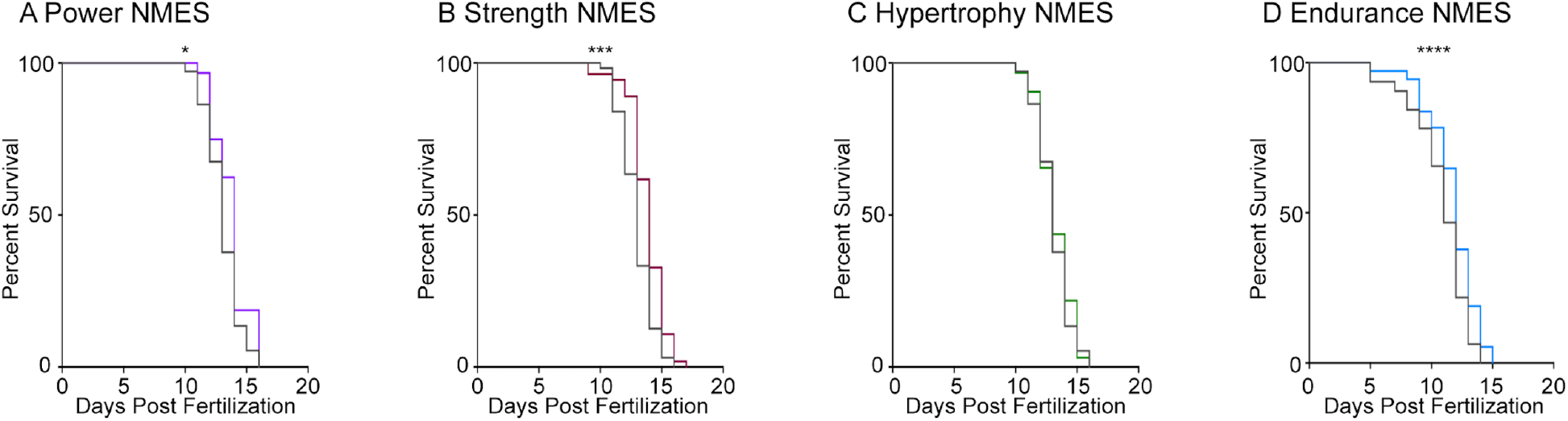
Changes in muscle health and swim activity do not predict survival. Survival was tracked following completion of the three NMES sessions. Survival was significantly improved in *dmd* mutants that completed power (A), strength (B), and endurance (C) NMES. (D) Hypertrophy NMES had no effect on survival in *dmd* mutants. Survival data were analyzed using a Mantel-Cox test. * p < 0.05, *** p < 0.001, **** p < 0.0001.

### eNMES improves muscle structure and sarcomere length

We found that eNMES significantly reduced muscle fiber detachments in *dmd* mutants (n = 22 control, n = 18 eNMES; p = 0.0191) (Figure 6F2). Many of the muscle segments in control *dmd* mutants had disorganized muscle fibers with a characteristic “waviness” and this appeared improved with eNMES. In order to quantify this aspect of muscle health, we used machine learning and trained the computer to identify, pixel-by-pixel, ‘healthy’ versus ‘sick’ with 97% accuracy. Next, we asked the computer to identify the percentage of healthy muscle in the same phalloidin images in which fiber detachments were counted on. From this analysis, we observed that *dmd* mutants completing three sessions of eNMES trend towards having higher percentages of health muscle compared to control *dmd* mutants (Figure 9A).

**Figure 9:**
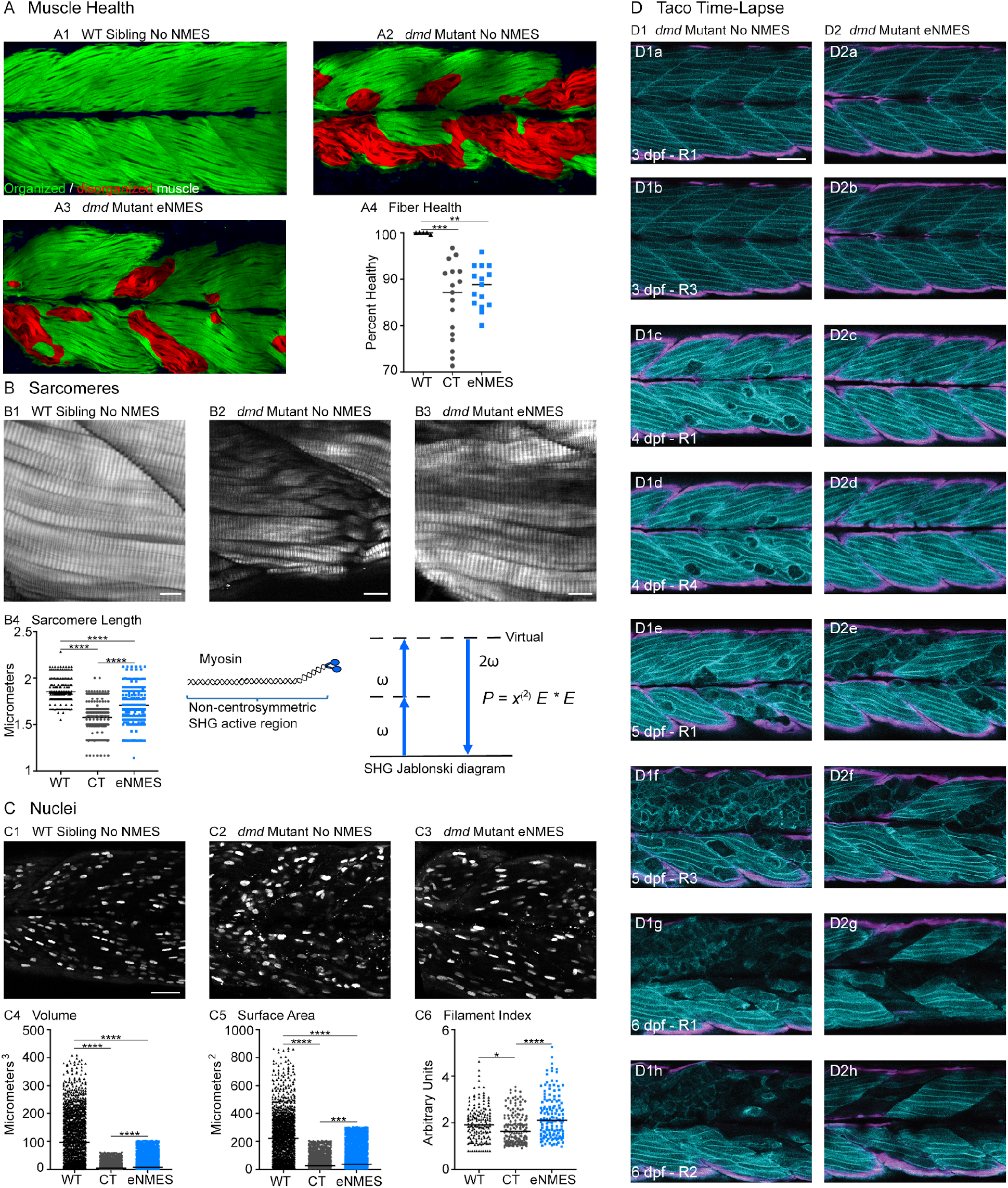
eNMES improves multiple components of muscle health. (A) Machine learning was used to quantify muscle health pixel-by-pixel. Green indicates healthy pixels while red indicates unhealthy pixels. (B) Second harmonic generation microscopy was used to quantify sarcomere length at 8 dpf. Representative SHG images of WT sibling control (B1), *dmd* mutant controls (B2), and *dmd* mutants that completed eNMES training. Anterior left, dorsal top, side mounted. Scale bars are 10 micrometers. (B4) Sarcomere length is significantly shorter in *dmd* mutant controls compared to WT sibling controls. However, eNMES significantly improves sarcomere length, bringing it closer to WT lengths. Each point represents a single sarcomere along a predetermined length of a muscle fiber. Multiple muscle fibers were measured per zebrafish. (C) Muscle nuclei were imaged at 8 dpf as a potential mechanism for improved muscle health. Anterior left, dorsal top, side mounted. (C1) Representative image of WT sibling control demonstrates healthy ellipsoidal nuclei organized along the length of the muscle fibers. (C2) Representative image of *dmd* mutant control demonstrates fragmented punctae as well as more spherical nuclei that clustering within the muscle segments. (C3) Representative image of *dmd* mutant that completed eNMES training demonstrates healthier, ellipsoidal nuclei that appear more organized within the muscle segments. Quantification of nuclear size indicates that eNMES significantly increases the volume (C4) and surface area (C5) of muscle nuclei compared to *dmd* mutant controls. However, nuclei are still significantly smaller compared to WT sibling controls. visually appear to have an increased number of myonuclei compared to unstimulated *dmd* mutants. (C6) Filament index was used to assess circularity, specifically the departure from a circle. Filament index is significantly higher in *dmd* mutants that completed eNMES training, indicating that nuclei are more elongated compared to *dmd* mutant controls. Each point represents a single nuclei within a z-stack. (D) Transgenic *dmd* mutants (mylpfa:lyn-cyan (cyan), smych1:GFP (magenta) were used to visualize changes in structural integrity of fast- and slow-twitch muscle fibers across three days. Anterior left, dorsal top, side mounted. Scale bar is 50 micrometers. Images were taken around the 12th myotome. (D1) Representative *dmd* mutant control. (D1a – D1b) At 3 dpf, there is no dystrophy in the imaged myotomes. (D1c – D1e) At 4 dpf and the beginning of 5 dpf, dystrophy is minimal with relatively few detaching muscle fibers. (D1f) However, massive muscle degeneration occurs between the first found of imaging and the third round of imaging at 5 dpf. (D1g-D1h) Fiber degeneration is still present, suggesting that the damaged muscle fibers have not been cleared and regeneration is unlikely. (D2) Representative *dmd* mutant that is undergoing eNMES training. (D2a - D2b) The first session of eNMES at 3 dpf does not result in immediate damage to the muscle. (D2c - D2d) Similarly, following the second session of eNMES at 4 dpf, there is no immediate muscle damage occurring in the imaged myotomes. (D2e) At 5 dpf, following the third session of eNMES, muscle fiber degeneration is evident but by the third round of imaging (D2f), these damaged areas are being cleared and there is evidence of regeneration. (D2g - D2h) At 6 dpf, previously damaged muscle segments have new muscle fibers present. All data were analyzed using either an ordinary one-way ANOVA with Tukey’s multiple comparisons test or a Kruskal-Wallis test with Dunn’s multiple-comparison test.* p < 0.05, ** p < 0.01, *** p < 0.001, **** p < 0.0001.

Sarcomere length impacts muscle function (Moo and Herzog, 2018). Sarcomeres produce force through the cross-bridges formed between actin and myosin, and the amount of force generated is dependent upon the amount of overlap between these thick and thin filaments (Gordon et al., 1966). We used Second Harmonic Generation (SHG) microscopy to investigate sarcomere structure at 8 dpf. SHG is a nonlinear process in which 2 photons of frequency ω designated as (E *E) interact with the non-centrosymmetric aligned dipole (χ^(2)^) region of the myosin tail. A single photon with twice the frequency (2 ω) and half the wavelength is emitted in this energy conserving label-free process. WT siblings exhibited a mean sarcomere length of 1.853 ± 0.1071 micrometers (Figure 9B4). This result corresponds with sarcomere lengths previously published in 3 dpf WT zebrafish (1.86 ± 0.15 micrometers; (Huang et al., 2011)). Sarcomeres are significantly shorter in *dmd* larvae (n = 356 sarcomeres *dmd* control, n = 282 sarcomeres WT sibling; p < 0.0001), with a mean length of 1.575 ± 0.1567 micrometers (Figure 9B4). This result corresponds with previous studies that showed shorter sarcomeres in *dmd* larvae (Widrick et al., 2016). We tested the hypothesis that eNMES would increase sarcomere length. Three sessions of eNMES significantly increased mean sarcomere lengths (1.707 ± 0.1710 micrometers) compared to *dmd* mutant controls (n = 356 sarcomeres *dmd* control, n = 550 sarcomeres eNMES; p < 0.0001) (Figure 9B4). Although eNMES-treated *dmd* larvae still have shorter sarcomeres than WT siblings, these data suggest the hypothesis that eNMES may improve muscle structure and function by restoring sarcomere lengths to more optimal lengths.

### Muscle nuclei return to a more ellipsoidal shape with eNMES

Myonuclear size and shape play an important role in muscle health (Folker and Baylies, 2013; Roman and Gomes, 2018). We measured three components of muscle nuclei size and shape: volume, surface area, and filament index. Filament index is a measure that quantifies the departure of an object from a circle. A circle has a filament index of 1 and a higher filament index indicates a departure to a more ellipsoidal shape. Thus, a higher filament index indicates that nuclei are more elongated, which is suggested to be healthier (Bruusgaard et al., 2003). Muscle nuclei in *dmd* mutants have significantly lower volumes (n = 1417 nuclei *dmd* control, n = 1355 nuclei WT sibling; p < 0.0001), surface areas (n = 1451 nuclei *dmd* control, n = 1378 nuclei WT sibling; p < 0.0001), and filament indices (n = 156 nuclei *dmd* control, n = 158 nuclei WT sibling; p = 0.0187) compared to WT siblings (Figure 9C4-6). Interestingly, eNMES significantly increased these measures (volume: n = 1417 nuclei *dmd* control, n = 1861 nuclei eNMES, p < 0.0001; surface area: n = 1451 nuclei *dmd* control, n = 1990 nuclei eNMES, p < 0.0001), especially for filament index (n = 156 nuclei *dmd* control, n = 140 nuclei eNMES; p < 0.0001), which is restored to WT values (Figure 9C6). Additionally, these nuclei appear more organized along the length of individual muscle fibers (Figure 9C3), similar to the pattern observed in WT siblings. These data suggest that *dmd* mutants have smaller, spheroidal nuclei compared to WT siblings, and eNMES is capable of elongating the nuclei, and increasing their volumes and surface areas.

### Longitudinal confocal analysis suggests less degeneration and more regeneration with eNMES

We used transgenic zebrafish (a generous gift from Drs. Sharon Amacher and Jared Talbot; (Hromowyk et al., 2020)) to visualize muscle structure through time. Disease onset in these transgenic zebrafish is at 3 dpf; thus, NMES sessions are at 3, 4 and 5 dpf while the recovery period extends from 6 through 9 dpf. At 3 dpf, there was not a clear difference in muscle degeneration between treated and control mutants. However, by 4 dpf, control mutants exhibited initial signs of muscle degeneration (Figure 9D1c and D1d). eNMES mutants showed less degeneration, suggesting that eNMES delays degeneration (Figure 9D2c and D2d). Whereas degenerated fibers persist in control mutants for days (Figure 9D1f - g), degenerated segments are cleared more quickly in eNMES-treated mutants (Figure 9D2f - g). Finally, more robust regeneration was observed in eNMES-treated mutants (Figure 9D2h). Taken together, these data suggest that eNMES improves muscle homeostasis.

### *Dmd* mutants respond differently than WT controls to eNMES

RNAseq was conducted on RNA extracted at 7 dpf, three days following the last NMES session. PCA analysis revealed that WT siblings that underwent three sessions of eNMES cluster separately from those in the control group (data not shown), suggesting that WT siblings that complete three sessions of eNMES have an unique expression profile compared to WT controls. Of the total 25,863 genes identified across the RNAseq analysis, 932 genes were differentially expressed between WT siblings in the eNMES versus control groups (Figure 10A1). Of these 932 genes, 306 genes were increased and 626 genes were decreased (Figure 10A3). Twenty-four GO terms were identified, including regulation of metabolic processes, regulation of MAP kinase activity, regulation of transcription, and circadian rhythms.

**Figure 10:**
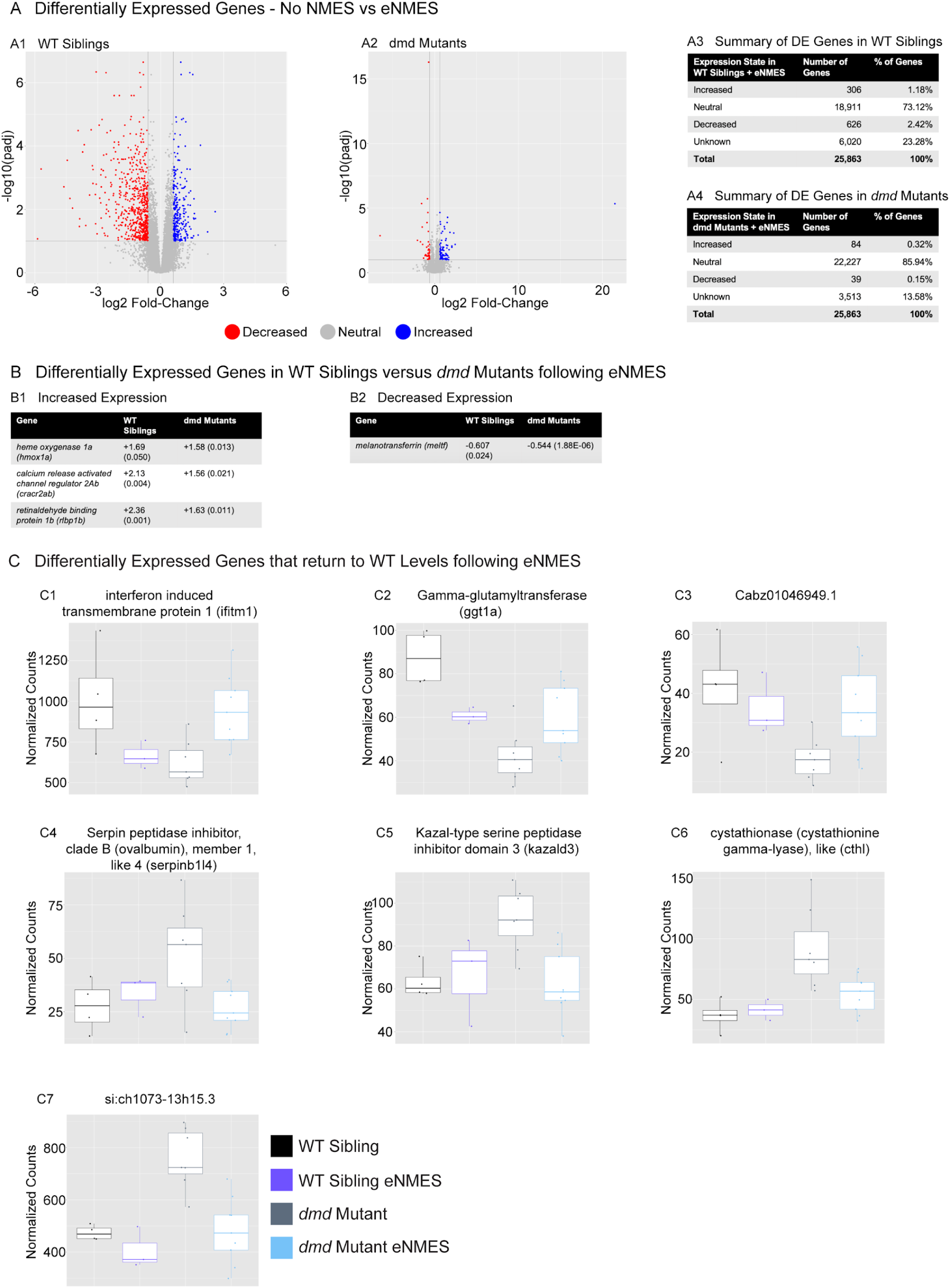
*dmd* mutants do not respond to eNMES in the same manner as WT siblings. RNAseq analysis was performed at 7 dpf in WT siblings and *dmd* mutants that completed eNMES training and their expression patterns were compared with their respective controls. (A1) Volcano plot showing significantly and biologically upregulated (blue dots) and downregulated (red dots) genes in WT siblings that completed eNMES versus those that did not. (A2) Volcano plot showing significantly and biologically upregulated (blue dots) and downregulated (red dogs) genes in *dmd* mutants that completed eNMES versus those that did not. (A3 – A4) Summary of differentially expressed genes in WT siblings (A3) and *dmd* mutants (A4). WT siblings had 932 differentially expressed genes compared to 123 differentially expressed genes in *dmd* mutants, suggesting that *dmd* muscle responds differently to eNMES and the genes responsible for eliciting beneficial effects on muscle structure and function are different. (B) Differentially expressed genes in WT siblings versus *dmd* mutants following eNMES that increased in expression (B1) or decreased in expression (B2). Values are presented as Fold Change (FDR). The relatively few genes that were significantly differentially expressed in both data sets further suggests that *dmd* mutants respond differently to eNMES than WT siblings. (C) Selected differentially expressed genes in *dmd* mutants following eNMES to show how expression levels return to those of WT controls.

Interestingly, the differentially expressed genes suggest that eNMES may be eliciting changes in WT versus *dmd* mutants through different mechanisms. Based on the number of genes differentially expressed, *dmd* mutants do not respond to NMES in the same manner as WT siblings (Fig. 10A2). One hundred and twenty-three genes were differentially expressed (FDR < 0.1 and abs(log2(Fold Change)) > 0.6) between *dmd* mutants that completed three eNMES sessions versus *dmd* mutant controls (Fig. 10A4). This number is much lower than the 932 differentially expressed genes in WT siblings. Additionally, *dmd* mutants have more genes that were increased (n = 84) than decreased (n = 39), which is the opposite of WT siblings, further suggesting that *dmd* mutants do not respond through the same signaling pathways as their healthy counterparts. Unfortunately, GO analyses did not reveal specific cellular processes in which these genes may participate in to positively impact muscle health. Only 4 genes of the 1,048 differentially expressed genes elicited by eNMES in both *dmd* mutants and WT siblings shared the same expression pattern (Fig. 10B). Three genes, including heme oxygenase 1a (*hmox1a*), calcium release activated channel regulator 2Ab (*cracr2ab*), retinaldehyde binding protein 1b (*rlbp1b*), were increased and 1 gene, melanotransferrin (*meltf*) was decreased with eNMES. This lack of overlap between differentially expressed genes further indicates that *dmd* mutants do not respond similarly to eNMES as WT siblings.

### eNMES may bring *dmd* mutants closer to WT transcription levels

We found 3 genes that exhibited opposite expression between genotypes (*dmd* mutants versus WT siblings) and eNMES (*dmd* mutants versus WT siblings after eNMES) (Fig. 10C). These genes were decreased in *dmd* mutants compared to WT siblings, but eNMES increased expression in *dmd* mutants and decreased expression in WT siblings. These genes included interferon induced transmembrane protein 1 (*ifitm1*), gamma-glutamyltransferase 1a (*ggt1a*), and Cabz01046949.1 (Figure 10C1-3). Additionally, 3 genes were increased in *dmd* mutants compared to WT siblings, but eNMES decreased expression in *dmd* mutants and had no effect in WT siblings. These genes included serpin peptidase inhibitor, clade B (ovalbumin), member 1, like 4 (*serpinb1l4*), kazal-type serine peptidase inhibitor domain 3 (*kazald3*), and cystathionase (cystathionine gamma-lyase), like (*cthl*) (Fig. 10C4-6). Lastly, 1 gene, si:ch107313h15.3, was increased in *dmd* mutants compared to WT siblings and was decreased in both WT siblings and *dmd* mutants following eNMES (Fig. 10C7). Altogether, these data suggest the possibility that eNMES may partially improve dmd muscle structure and function by returning *dmd* mutants to more WT-like expression profiles.

### eNMES may alter the ECM in *dmd* mutants

The ECM surrounding muscle fibers is a critical component of muscle fiber health. Protein complexes spanning the sarcolemma and ECM serve as mechanical linkages and signaling hubs that promote muscle plasticity (Csapo et al., 2020). However, excess ECM protein deposition can also lead to fibrosis. We asked whether ECM proteins are differentially expressed in zebrafish *dmd* larvae following eNMES. Despite the fact that RNAseq data represent a snapshot in time and are not the best way to capture a structure as dynamic as the ECM, we observed changes in ECM gene expression with eNMES.

Transforming growth factor beta induced (TGFBI) is an extracellular matrix protein that binds to type I, II and IV collagens as well as several integrins. Tgfbi is upregulated in *mdx* muscle compared to healthy muscle (Coles et al., 2020; Pescatori et al., 2007). We found that *tgfbi* is also significantly higher in zebrafish *dmd* mutants compared to WT controls. Expression of *tgfbi* in both *dmd* mutants and WT siblings was reduced with eNMES (Figure 11A1). Periostin (postnb) is a TGFBI-related protein that is involved in modeling the ECM and connective tissue architecture during development and regeneration, serving specifically as a mediator of fibrosis in injury and disease (Ozyilmaz et al., 2019). RNAseq data indicate that *postnb* shares a similar expression pattern with *tgfbi*: increased expression in *dmd* mutants compared to WT siblings and a reduction in this expression following eNMES in both groups (Figure 11A2). Integrin-β1 (*itgb1b.2*) is also significantly upregulated in *dmd* mutants compared to WT siblings, but is reduced with eNMES (Figure 11A3). In skeletal muscle, integrins are a family of cell surface adhesion molecules that mediate cell-matrix interactions. Altogether, these data suggest that cell-ECM interactions and the composition of the ECM are changed with eNMES.

**Figure 11:**
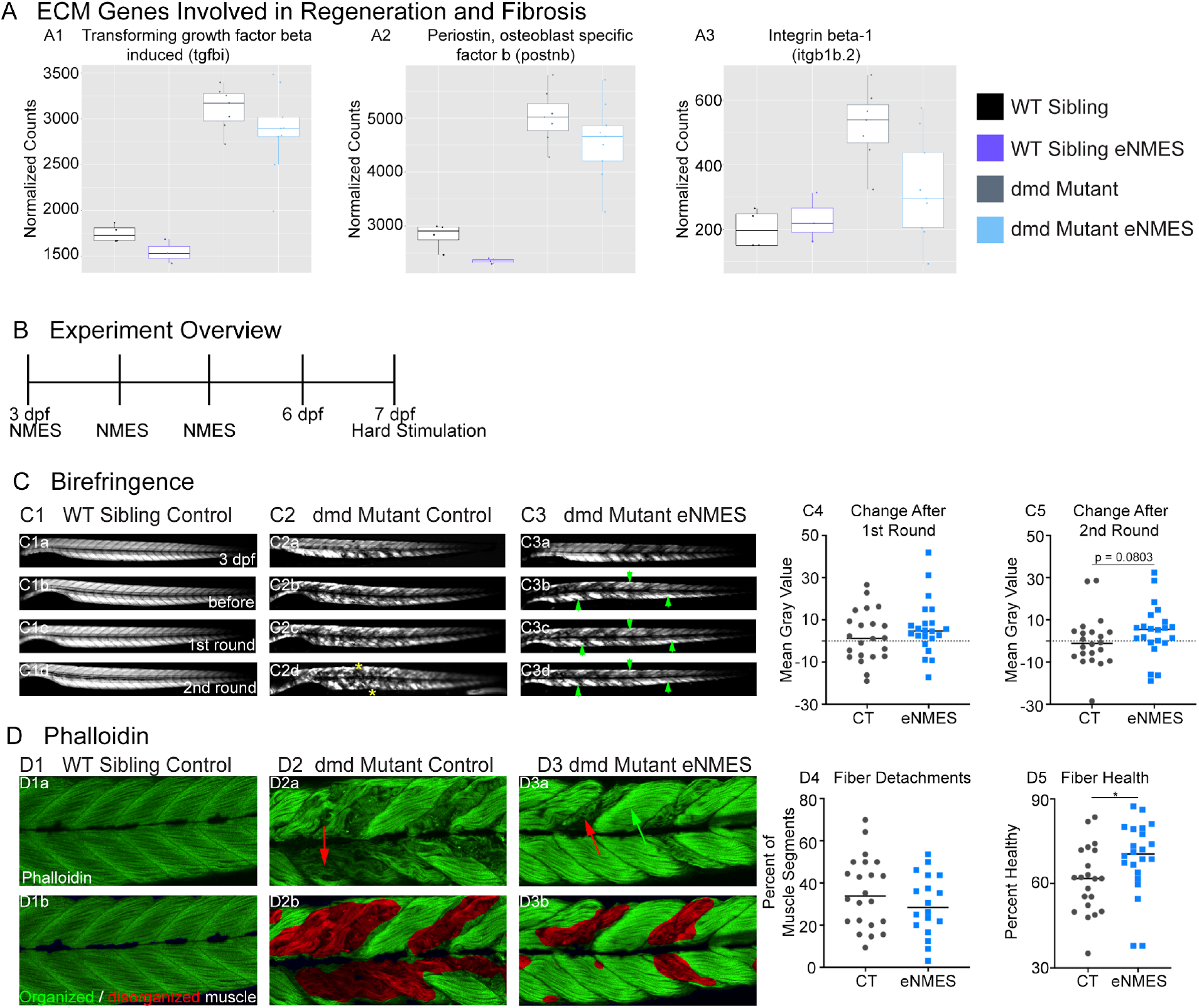
Modulation of ECM genes involved in regeneration and fibrosis following eNMES may lead to observed improvements in muscle resilience in *dmd* mutants. We identified three ECM genes, tgfbi (A1), postnb (A2), itgb1b.2 (A3) that are significantly upregulated in *dmd* mutants compared to WT siblings and trend to be downregulated with eNMES in *dmd* mutants. (B) Experiment overview. At 3 dpf (disease onset), birefringence images are taken followed by the first session of eNMES. At 4 and 5 dpf, zebrafish undergo the second and third NMES sessions, respectively. At 7 dpf, muscle resilience was tested using a hard electrical stimulation paradigm intended to cause muscle damage. (C) Birefringence images were taken at 3 dpf, before the first session and after the first and second sessions. (C1) No visible changes in birefringence are observed in WT siblings after the two stimulation sessions. (C2) For *dmd* mutant controls, the first round of stimulation did not result in visible changes to birefringence (C2c) but, after the second round, areas of muscle degeneration are visible (C2d, yellow asterisks). Conversely, in *dmd* mutants that completed three sessions of eNMES, the first (C3c) and second (C3d) rounds of stimulation did not result in visible changes to birefringence (green arrowheads denote intact areas of birefringence that remain intact). (C4, C5) Change in birefringence from before to after the first round (C4) and second (C5) of stimulation suggests that eNMES training may improve muscle resilience. (D) Phalloidin was used to visualize individual muscle fibers. (D1a) Representative image of a WT sibling control demonstrates healthy, organized muscle fibers and myotomes. (D2a) Representative image of a *dmd* mutant control highlights disorganized and wavy muscle fibers and fiber detachments. (D3a) Representative image of a *dmd* mutant that completed eNMES demonstrates some wavy muscle fibers and detached fibers intermixed with relatively healthy myotomes. (D4) The percent of muscle segments with detached fibers following the hard stimulation is reduced in *dmd* mutants that complete eNMES training compared to *dmd* mutant controls. For this analysis a muscle segment was defined as half of a myotome. (D1b, D2b, D3b) Machine learning was used to quantify muscle health pixel-by-pixel. Green indicates healthy pixels while red indicates unhealthy pixels. (D5) The percent of healthy muscle following the hard stimulation is significantly higher in *dmd* mutants that completed eNMES compared to *dmd* mutant controls. All data were analyzed using two-sided t tests. * p < 0.05.

### eNMES may reduce susceptibility to contraction-induced injury in *dmd* mutants

Cell-matrix adhesion is negatively affected in various models of muscular dystrophy and restoration of adhesion improves muscle structure and function (Burkin et al., 2005, 2001; Goody et al., 2012). Therefore, the downregulation of key cell adhesion proteins following eNMES was puzzling and led us to ask whether muscle cell-matrix adhesion was altered by eNMES. We did this by subjecting zebrafish to a hard stimulation paradigm designed to make muscle fibers detach from their ECM for two back-to-back sessions (Figure 11B). This experiment was conducted 2 days after the final eNMES training session. This one minute hard stimulation paradigm was defined by a frequency of 4 pulses per second, a delay of 60 ms, a duration of 2 ms, and a voltage of 30 volts, which is similar to that known to initiate muscle fiber detachment from their ECM (Subramanian and Schilling, 2014). Birefringence images were taken before and after each session. To ensure consistency in imaging, zebrafish were mounted laterally with their left side facing up, and the same imaging parameters were used for each zebrafish across all imaging sessions. We then analyzed the change in mean gray values before stimulation compared to after the first or second session. Nearly half of control mutants (10/22) had decreased mean gray values after the first session (Fig. 11C4), and slightly over half (13/22) had decreased mean gray values after the second session (Fig. 11C5). In contrast, just under 25% of eNMES treated mutants (5/22) had a decreased mean gray value after the first session (Fig. 11C4) and slightly under a third had a decreased mean gray value after the second session (7/22; Fig. 11C5). While there are no differences in absolute mean gray values between control and eNMES mutants before and after the first round of stimulation (Fig. 11C4), the change in mean gray values for eNMES-treated *dmd* mutants is higher than controls following the second round of stimulation (Fig. 11C5). These birefringence data suggest that eNMES may improve cell-adhesion. If this is the case, we would predict that there would be fewer muscle detachments in eNMES mutants after a hard stimulation compared to control mutants after a hard stimulation. Indeed, the percent of muscle segments with detachments trends lower in *dmd* mutants that completed eNMES compared to control *dmd* mutants (Figure 11D4). The most striking difference in appearance between the control and eNMES *dmd* mutants after the hard stimulation was the improved organization of muscle fibers in eNMES treated *dmd* mutants. Whereas control *dmd* mutants had lots of disorganized fibers (red arrow, Fig. 11D2a), eNMES *dmd* mutants had more organized fibers (green arrow, Fig. 11D3a). We used machine learning to quantify overall muscle health. This approach showed that *dmd* mutants that completed eNMES had a significantly higher percentage of healthy muscle compared to control *dmd* mutants (n = 21 control, n = 22 eNMES; p = 0.0496) (Figure 11D5). Taken together, these data indicate that eNMES-treated *dmd* mutants can withstand contraction-induced injury better than *dmd* mutant controls.

## DISCUSSION

We used an experimental design that leverages the power of the zebrafish model’s ability to perform in-vivo analyses of numerous components of organismal health across time in individual zebrafish. By implementing this longitudinal design, we demonstrate that (1) the zebrafish model for DMD exhibits phenotypic variation in disease progression; (2) periods of inactivity are detrimental to DMD muscle health and survival; (3) endurance NMES positively benefits muscle health, function, and survival in *dmd* mutants; (4) changes are accompanied by improvements in NMJ length, nuclear shape and size, and sarcomere lengths; and, (5) *dmd* mutants respond to NMES differently than WT siblings. These findings indicate that the zebrafish model is a valuable tool for studying skeletal muscle plasticity and that healthy and dystrophin-deficient muscle use different mechanisms to maintain homeostasis.

### Phenotypic variation in the zebrafish model of *dmd*

The clinical presentation of muscular dystrophies is frequently variable: ranging from severe, congenital muscle weakness to mild, adult-onset muscle weakness. Similarly, variability across individuals with the same disease-causing allele is common. This variability makes it difficult to accurately inform patients as to how their disease will progress and/or respond to therapies. One roadblock to understanding the phenotypic spectrum of muscular dystrophies is that the basic biological mechanisms of variability in musculoskeletal development and disease are not well understood. To our knowledge, we are the first to demonstrate that *dmd* mutant zebrafish exhibit significant variation in disease severity. The mild phenotype involves very few muscle segments with visible signs of dystrophy at the onset of muscle degeneration, and the severe phenotype presents with many muscle segments with muscle fiber detachments and disorganized fibers. This variation at disease onset is extremely important to identify since disease progression, especially in the first three days after disease onset, is different between the two groups. During this time, mild *dmd* mutants are degenerating while severe *dmd* mutants are regenerating. Surprisingly, even though mild and severe mutants exhibit similar muscle structure at 8 dpf, their initial severity level continues to affect their gross muscle function. Severe *dmd* mutants swim a significantly lower distance and at a significantly slower mean velocity at 5 dpf and at 8 dpf compared to mild *dmd* mutants. Therefore, recognizing that there are differences in muscle homeostasis between mild and severe *dmd* mutants is critical for experimental design, especially to ensure that variables such as disease stage at time of treatment and at time of evaluation are not influencing outcome measures. In the future, studying variation in the context of differentially expressed genes could unveil potential biomarkers for neuromuscular plasticity and their potential roles in muscle structure, function and survival. Collectively, these data will be valuable when evaluating pharmacological interventions as some may have differential effects based on the current expression levels of the targeted pathways in mild versus severe *dmd* mutants.

### Long-term inactivity is deleterious in the fish model of *dmd*

A major consequence of muscle wasting and weakness caused by muscular dystrophy is dramatic reductions in physical activity levels. For example, step activity patterns from individuals with DMD indicate that activity levels are lower (McDonald et al., 2005). While this loss in activity is a clear consequence of muscle wasting in the absence of dystrophin, the consequences of this reduced activity on advancing muscle wasting in DMD have not been extensively studied. Inactivity protects muscle from damage (Berger et al., 2010; Mizuno, 1992; Mokhtarian et al., 1999), and our extended inactivity data confirm that muscle structure is protected *during the inactive period*. However, the consequences of resumption of activity have not been well investigated. To our knowledge, only one study examined how inactive muscle fibers respond to the reinstatement of activity. In *mdx* mice from Hourdé et al.’s (2013) study and in our *dmd* mutant zebrafish, this protective effect is quickly reversed when inactive muscle is subjected to activity. Data from both studies demonstrate that inactive muscle is more susceptible to damage compared to freely moving controls. Therefore, these data clearly demonstrate that inactivity worsens disease progression in animal models of dystrophin-deficient muscle. Future studies should evaluate the long-term consequences of prolonged reductions in activity in individuals with DMD because muscle resiliency has major impacts on disease progression.

### Zebrafish as a model for elucidating neuromuscular plasticity

The negative consequences of inactivity on muscle resilience led us to ask whether activity could improve muscle resilience, and, therefore, disease progression. We selected NMES as a mechanism to elicit consistent, repeatable contraction patterns across individual zebrafish. We generated four NMES paradigms that varied in frequency and voltage to test how different contraction patterns impact muscle structure, function and survival. Collectively, our experiments suggest that *dmd* muscle exhibits a delicate, intricate equilibrium with several factors influencing muscle structure, swimming activity and survival. Birefringence does not predict swimming performance and swimming performance does not predict survival. For example, two paradigms, eNMES and pNMES, improved muscle structure while two paradigms, hNMES and sNMES, negatively affected muscle structure. Surprisingly, though, only eNMES increased swimming activity. In contrast, survival was extended by eNMES as well as pNMES and sNMES. Therefore, this is a new model to understand disease progression and elucidate mechanistic pathways that target improvements in structure, function, and survival.

### Potential mechanisms for improved neuromuscular function

We found that eNMES increased sarcomere lengths, which could improve force generation by leading to a more optimal interaction between actin and myosin filaments. Nuclear volume, surface area, and filament index were increased with eNMES, suggesting that muscle nuclei are returning to their elongated shape. As nuclear size affects DNA organization, transcriptional and translational processes, and nuclear import and export activities (Levy and Heald, 2012), minor changes in size correlate with reduced muscle function and fiber performance (Windner et al., 2019). Therefore, these improvements in muscle nuclei may also mediate improvements in muscle structure and function following eNMES. Time-lapse imaging data support the hypothesis that eNMES is creating an environment that supports regeneration. Following eNMES in *dmd* mutants, there is less degeneration and in those areas with degenerating fibers, newly regenerated fibers appear sooner. Lastly, RNAseq data identify two potential mechanisms that may allow for the above improvements to occur. The first mechanism includes restoring gene expression levels back to those in WT siblings. In several genes, we see expression levels increasing or decreasing towards those levels of WT sibling controls following eNMES in *dmd* mutants. More often than not, these expression changes are opposite what is seen in WT siblings that completed eNMES. The second mechanism includes potential remodeling of the ECM. The ECM is constantly responding to signals from both within and outside the cell, and incorporating these signals to create a scaffold that supports either regeneration or fibrosis such that the cell is protected from further damage. Our RNAseq data suggest that eNMES may result in ECM remodeling to support regeneration and/or limit fibrosis.

### Summary

Identifying the basic mechanisms by which inactivity and activity impact muscle health in the context of muscle disease is a crucial first step towards identifying potential therapies. Here, we show that inactivity is deleterious in the zebrafish model of DMD. We also identify a NMES paradigm that improves neuromuscular structure, function, and lifespan. Significantly, we show that NMES differently affects gene expression in WT versus *dmd* mutants. This result indicates that it is critical to study the impacts of activity on diseased muscle in addition to WT muscle. Taken together, our data not only establish a model system for neuromuscular plasticity in healthy versus diseased muscle; but also clearly elucidate deleterious effects of inactivity and beneficial effects of NMES.

## Acknowledgements

The authors would like to thank Drs. Sharon Amacher and Jared Talbot for developing the transgenic 3MuscleGlow zebrafish and sharing this valuable tool with us; Dr. Joy-El Talbot at Iris Data Solutions for her expertise in RNAseq analysis; NVIDIA Corporation for donating the Quadro P6000 used for deep learning analyses; Keegan Kilroy for designing the NMES gym and assistance with designing the NMES paradigms; and Mark Nilan for exceptional zebrafish care at the UMaine Zebrafish Facility.

